# Rad51 paralog complex Rad55–Rad57 acts as a molecular chaperone during homologous recombination

**DOI:** 10.1101/2020.11.04.367136

**Authors:** Upasana Roy, Youngho Kwon, Lea Marie, Lorraine Symington, Patrick Sung, Michael Lisby, Eric C. Greene

## Abstract

Homologous recombination (HR) is essential for the maintenance of genome integrity. Rad51 paralogs fulfill a conserved, but undefined role in HR, and their mutations are associated with increased cancer risk in humans. Here, we use single–molecule imaging to reveal that the *Saccharomyces cerevisiae* Rad51 paralog complex Rad55–Rad57 promotes the assembly of Rad51 recombinase filaments through transient interactions, providing evidence that it acts as a classical molecular chaperone. Srs2 is an ATP–dependent anti–recombinase that downregulates HR by actively dismantling Rad51 filaments. Contrary to the current model, we find that Rad55– Rad57 does not physically block the movement of Srs2. Instead, Rad55–Rad57 promotes rapid re– assembly of Rad51 filaments after their disruption by Srs2. Our findings support a model in which Rad51 is in flux between free and ssDNA–bound states, the rate of which is dynamically controlled though the opposing actions of Rad55–Rad57 and Srs2.

## Introduction

Homologous recombination (HR) is crucial for the repair of stalled or collapsed replication forks, DNA double–strand breaks (DSB), and chromosome segregation during meiosis (Kowalczykowski, 2015; Mehta and Haber, 2014; San Filippo et al., 2008). Lack of, or inappropriate HR, can lead to gross chromosomal rearrangements which are a hallmark of cancer cells (Kowalczykowski, 2015; Mehta and Haber, 2014; Prakash et al., 2015; San Filippo et al., 2008). The initial steps in HR involve extensive DNA resection around the DSB site to expose 3’ single–stranded DNA (ssDNA) ends that are rapidly bound by replication protein A (RPA) (Bonetti et al., 2018; Cejka, 2015; Symington, 2016). The key DNA transactions that occur during HR are catalyzed by members of the ATP–dependent Rad51/RecA family of DNA recombinases, which form extended helical filaments on the ssDNA associated with processed DNA breaks (Bianco et al., 1998; Kowalczykowski, 2015; Mehta and Haber, 2014; San Filippo et al., 2008). Once assembled, the Rad51–ssDNA filaments search for a homologous target in the genome, invade the DNA target to pair the broken DNA end with the latter to produce a DNA joint called the displacement–loop (D–loop). The broken DNA end is then extended by a DNA polymerase using the newly annealed homologous template, and resulting DNA intermediates can be resolved into either crossover or non–crossover products (Kowalczykowski, 2015; Mehta and Haber, 2014; San Filippo et al., 2008).

Given their central importance to HR and genome integrity, Rad51 recombinase filaments are targets of numerous positive and negative regulatory factors. Importantly, RPA inhibits Rad51– ssDNA filament assembly due to the high affinity of RPA for ssDNA. This inhibitory effect must be overcome with the help of positive HR regulators called “mediators” that stimulate Rad51 filament assembly on RPA–ssDNA (Bonilla et al., 2020; Kowalczykowski, 2015; Mehta and Haber, 2014; Prakash et al., 2015; San Filippo et al., 2008). Rad51 paralogs are classified as HR mediators and they are broadly conserved across eukaryotes, however, the mechanistic basis for Rad51 paralog function remains poorly defined (Bonilla et al., 2020; Prakash et al., 2015; Sullivan and Bernstein, 2018). Rad51 paralogs share ~20% identity to the conserved central ATPase core domain of Rad51, with little to no similarity outside this region (Bonilla et al., 2020; Prakash et al., 2015; Sullivan and Bernstein, 2018). There are five human RAD51 paralogs (RAD51B, RAD51C, RAD51D, XRCC2 and XRCC3), and cells lacking any of these proteins are sensitive to DNA damaging agents and exhibit chromosomal abnormalities (Bonilla et al., 2020; Prakash et al., 2015). Gene knockouts are embryonic lethal in mice (Prakash et al., 2015; Sullivan and Bernstein, 2018), and mutations in the human proteins have been identified in the cancer–prone syndrome Fanconi Anemia and a variety of cancers (Bonilla et al., 2020; Meindl et al., 2010; Pennington et al., 2014). Acting in opposition to RAD51 paralogs are “anti–recombinases”, such as RECQ5 and FBH1, which dismantle Rad51 filaments and prevent excessive or inappropriate recombination (Branzei and Foiani, 2007; Branzei and Szakal, 2017). Mutations in these anti–recombinases can also severely compromise genomic integrity and are associated with cancer (Branzei and Foiani, 2007; Branzei and Szakal, 2017). Despite their importance in human health, the regulatory interplay between RAD51 paralogs and anti–recombinases remains poorly understood.

The yeast Rad51 paralogs Rad55–Rad57 and the anti–recombinase Srs2 are important models for understanding positive and negative HR regulators. Srs2 is an ATP–dependent ssDNA motor protein that translocates on DNA in a 3’→ 5’ direction and strips Rad51 from the ssDNA (Antony et al., 2009; Kaniecki et al., 2017; Krejci et al., 2003; Rong and Klein, 1993; Veaute et al., 2003). Srs2 promotes dissociation of Rad51 from ssDNA in part by stimulating Rad51 ATP hydrolysis activity through a direct protein–protein interaction (Antony et al., 2009). Notably, the Srs2 interaction with Rad51 is important for its Rad51 disruption activity (Colavito et al., 2009), but it is not essential for Srs2 recruitment to HR foci (Burgess et al., 2009), suggesting that Srs2 may be recruited to recombination intermediates via other factors.

Rad55–Rad57 is a stable heterodimer *in vivo* and is approximately 10–fold less abundant than cellular Rad51 (Sung, 1997). Despite this low abundance, deletion of *RAD55* and *RAD57* sensitizes cells to ionizing radiation (IR), and this phenotype can be suppressed by Rad51 overexpression or by *SRS2* deletion (Fortin and Symington, 2002; Fung et al., 2009; Hays et al., 1995; Johnson and Symington, 1995). Although Rad55 and Rad57 contain the RecA/Rad51 ATPase core domain, they do not form filaments on ssDNA or carry out strand exchange themselves, and display only weak ATPase activity, that unlike Rad51, is not stimulated by DNA cofactors (Sung, 1997). Biochemical experiments show that Rad55–Rad57 promotes Rad51 filament assembly and strand exchange activity by overcoming the inhibitory effect of RPA, and not through a direct stimulation of Rad51 strand exchange activity, as reactions without RPA are not stimulated (Gaines et al., 2015; Sung, 1997).

Rad55–Rad57 and Srs2 share an antagonistic relationship, as *SRS2* deletion suppresses the IR sensitivity of *rad57* and *rad55* mutants, and in biochemical experiments Rad55–Rad57 suppresses Rad51–ssDNA disruption by Srs2 (Fung et al., 2009; Liu et al., 2011). A model has been put forth to suggest that Rad55–Rad57 associates stably with the Rad51–ssDNA nucleoprotein filament and physically impedes Srs2 translocation through a direct protein interaction with Srs2 to attenuate its anti–recombinase activity (Liu et al., 2011). Interestingly, the meiosis–specific recombinase Dmc1 is a potent inhibitor of Srs2 ATPase activity and highly adept at blocking Srs2 translocation (Crickard et al., 2018a; Sasanuma et al., 2013). In striking contrast to Dmc1, Rad55–Rad57 does not inhibit Srs2 ATP hydrolysis activity (Liu et al., 2011), indicating that it acts via a distinct, yet to be determined mechanism to counteract Srs2 activity.

Here, we define the role of Rad55–Rad57 as a recombination mediator and antagonist of Srs2, by visualizing Rad51–ssDNA dynamics in real–time at the single–molecule level. Importantly, in contrast to published work (Liu et al., 2011), we find that Rad55–Rad57 is not a stable co–component of the Rad51–ssDNA presynaptic complex, but instead associates with Rad51–ssDNA complexes during a transient burst that coincides only with the earliest stages of Rad51 filament assembly. The transient binding behavior of Rad55–Rad57 is reminiscent of a chaperone–like activity rather than the paralog complex serving as an integral component of the mature presynaptic complex. The dissociation of Rad55–Rad57 from the Rad51–ssDNA presynaptic complex requires ATP hydrolysis by Rad55. In addition, we show that Rad55–Rad57 stimulates both the rate and extent of RPA displacement by Rad51 but does not directly inhibit Srs2 motor activity. Instead, Rad55–Rad57 acts in opposition to Srs2 by facilitating rapid Rad51 filament re–assembly. Together, our findings lead to a new model to explain how the conserved Rad51 paralogs help maintain genome integrity through their regulation of the Rad51 filament assembly.

## Results

### Rad55–Rad57 does not interact with RPA–ssDNA complexes

We used ssDNA curtain assays with mCherry labeled RPA to observe Rad51 filament dynamics in the presence of Srs2 (De Tullio et al., 2018; De Tullio et al., 2017; Kaniecki et al., 2017); note, use of fluorescent RPA is necessary because fluorescent Rad51 fusion proteins are not fully functional (Lisby et al., 2004; Waterman et al., 2019). We generated an N–terminal GFP–tagged version of Rad55 for single molecule imaging of the Rad55–Rad57 complex using total internal reflection fluorescence microscopy (TIRFM; Figure 1A–C). GFP–tagged Rad55–Rad57 forms DNA repair foci *in vivo* in response to IR (Lisby et al., 2004). Expression of GFP–Rad55–Rad57 suppressed camptothecin sensitivity of *rad55Δ rad57Δ* strains, confirming that the GFP–tagged heterodimer complex assembles correctly and is functional (Figure S1A). We then purified GFP– Rad55–Rad57 complex as a stable heterodimer (Figure S1B), and affinity pull–down assays confirmed that the complex interacted with Srs2 (Figure S1C) (Liu et al., 2011).

**Figure 1.**
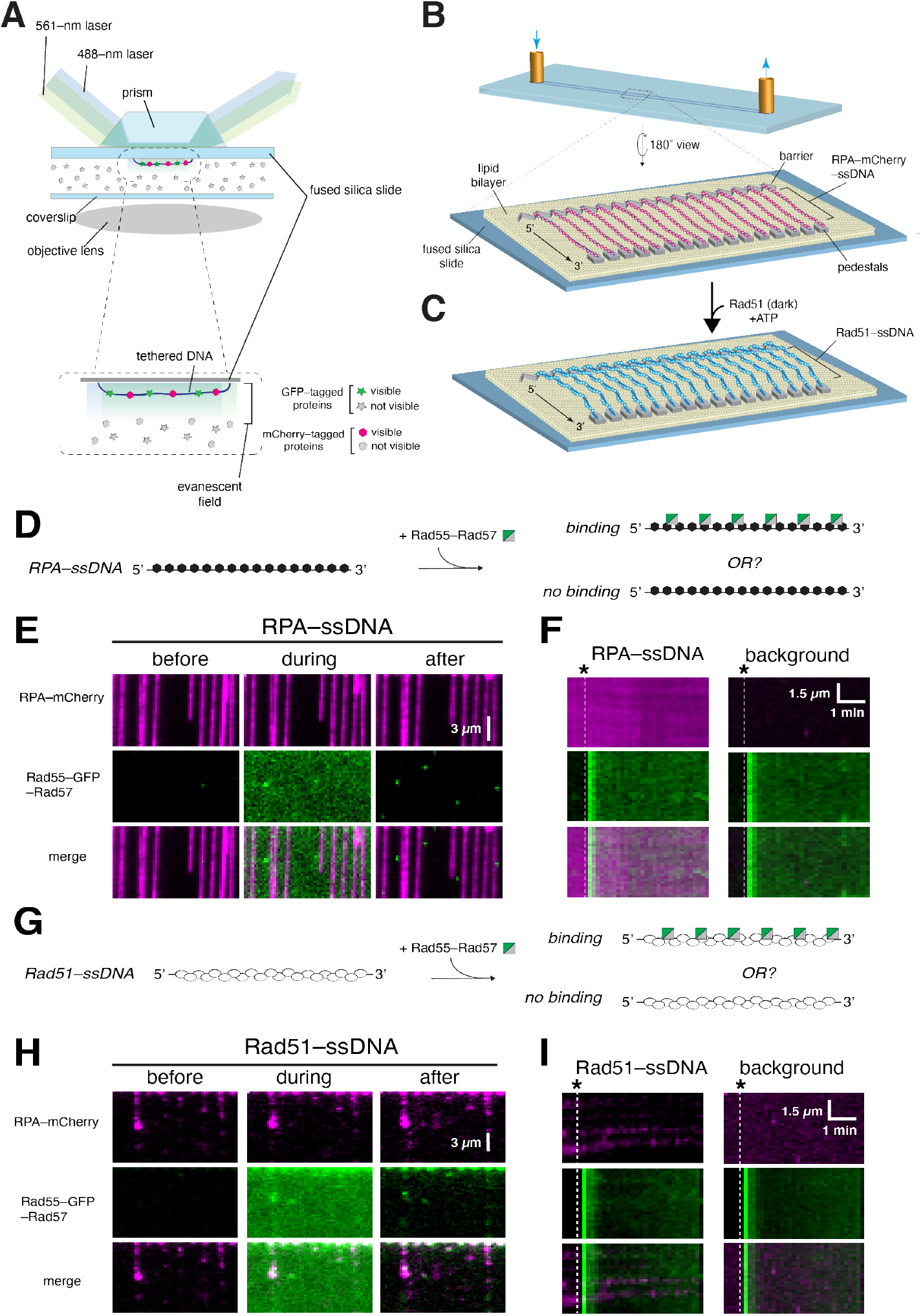
Rad55–Rad57 does not bind RPA–ssDNA or mature Rad51–ssDNA filaments. (**A**) Schematic illustration of two–color laser excitation with a prism–type TIRF microscope. (**B**), Flowcell and slide surface schematic showing the lipid bilayer, nanofabricated chromium barrier and pedestals, and tethered molecules of ssDNA bound by RPA–mCherry. (**C**) Schematic showing assembly of Rad51 filaments on the RPA–mCherry bound ssDNA. (**D**) Assay schematic to test binding of Rad55–Rad57 to RPA–ssDNA. (**E**) Wide–field view of RPA–mCherry (magenta) bound ssDNA 1 min before, during, and after injection of 60 nM GFP–tagged Rad55–Rad57 (green). (**F**) Representative kymographs of RPA–mCherry bound ssDNA, and a background region lacking ssDNA. Injection of GFP–Rad55–Rad57 is indicated by the white dashed line. (**G**) Assay schematic to test binding of Rad55–Rad57 to preassembled Rad51–ssDNA filaments. (**H**) Wide–field view of Rad51–ssDNA filaments (unlabeled) 1 min before, during, and after injection of 60 nM GFP–tagged Rad55–Rad57 (green). (**I**) Representative kymographs of Rad51–ssDNA and a background region lacking ssDNA. Injection of GFP–Rad55–Rad57 (green) is indicated by the white dashed line.

Rad55–Rad57 is classified as a mediator based on its ability to relieve inhibition by RPA on Rad51–mediated strand exchange (Sung, 1997). We therefore first examined Rad55–Rad57 interactions with RPA–ssDNA filaments, the physiologically relevant substrate for Rad51 assembly. We assembled ssDNA curtains bound with RPA–mCherry (De Tullio et al., 2018; Ma et al., 2017b), flushed out unbound RPA–mCherry from the sample chamber, and then injected GFP–Rad55–Rad57 in buffer containing 2 mM ATP (Figure 1D). Images were recorded every 10s over the course of the experiment (~20 mins) to visualize how Rad55–Rad57 interacts with the RPA–ssDNA complexes. Interestingly, Rad55–Rad57 did not bind the RPA–ssDNA complexes as shown in the wide–field images (Figure 1E) and kymographs of individual RPA–ssDNA molecules (Figure 1F). The finding that Rad55–Rad57 did not bind to RPA–ssDNA was consistent with the observation that Rad55–Rad57 foci do not form *in vivo* without Rad51 (Lisby et al., 2004).

### Rad55–Rad57 does not interact with mature Rad51 filaments

We next examined the interaction of Rad55–Rad57 with pre–assembled Rad51–ssDNA filaments. Rad51 was injected into a sample chamber containing ssDNA molecules bound by RPA–mCherry and incubated in the absence of buffer flow for 20 mins to allow Rad51–ssDNA filament assembly. Rad51–ssDNA assembly was monitored by the loss of mCherry–RPA signal from the ssDNA as previously described (De Tullio et al., 2018; Ma et al., 2017b). After this incubation period, unbound proteins were flushed from the sample chamber and GFP–Rad55–Rad57 was then injected while visualizing its interactions with the Rad51–ssDNA filaments. Surprisingly, we saw little to no binding of Rad55–Rad57 to pre–assembled Rad51 filaments (Figure 1G–I). This result was unexpected, given that Rad55–Rad57 interaction with Rad51 has been observed in both two–hybrid and immunoprecipitation studies (Godin et al., 2013; Hays et al., 1995; Johnson and Symington, 1995; Liu et al., 2011).

### Rad55–Rad57 binds transiently during Rad51 filament assembly

We considered the possibility that productive interaction between Rad55–Rad57 and Rad51 may require co–assembly of both onto ssDNA. To test this hypothesis, Rad51 and Rad55–Rad57 were co–injected into a sample chamber containing ssDNA molecules bound by RPA–mCherry in buffer supplemented with 2 mM ATP (Figure 2A). Under these conditions, there was extensive co–localization of Rad55–Rad57 and Rad51 on the ssDNA (Figure 2B), providing a satisfying explanation for the observation that Rad51 is required for Rad55–57 foci formation *in vivo* (Lisby et al., 2004). Remarkably, Rad55–Rad57 binding was highly transient, occurring only within a brief ~2 min window that coincided with the earliest phases of Rad51 filament assembly (Figure 2B–D). The vast majority of the Rad55–Rad57 complexes then quickly dissociated from the Rad51–ssDNA (*k*_off_ = 0.455 ± (0.423 – 0.488; 95% confidence interval (CI)) min^−1^; Figure 2D & Table S1), and within ~10 min the GFP–Rad55–Rad57 signal remaining on the mature Rad51– ssDNA filaments was indistinguishable from background (Figure S2). To further confirm our results, we imaged the Rad51–ssDNA after filament assembly had plateaued using maximum laser intensity (200mW) and found that 86% of ssDNA filaments had no detectable Rad55–Rad57 at these later time points (Figure 2E–G). Of the ssDNA molecules with residual Rad55–Rad57, most had only one detectable Rad55–Rad57 complex within an entire 36,000–nucleotide segment (Figure 2G). Together, our results show that Rad55–Rad57 binds to Rad51 filaments in a transient burst coinciding with the earliest stages of filament assembly and the vast majority then quickly dissociates from Rad51–ssDNA as filament assembly progresses towards completion.

**Figure 2.**
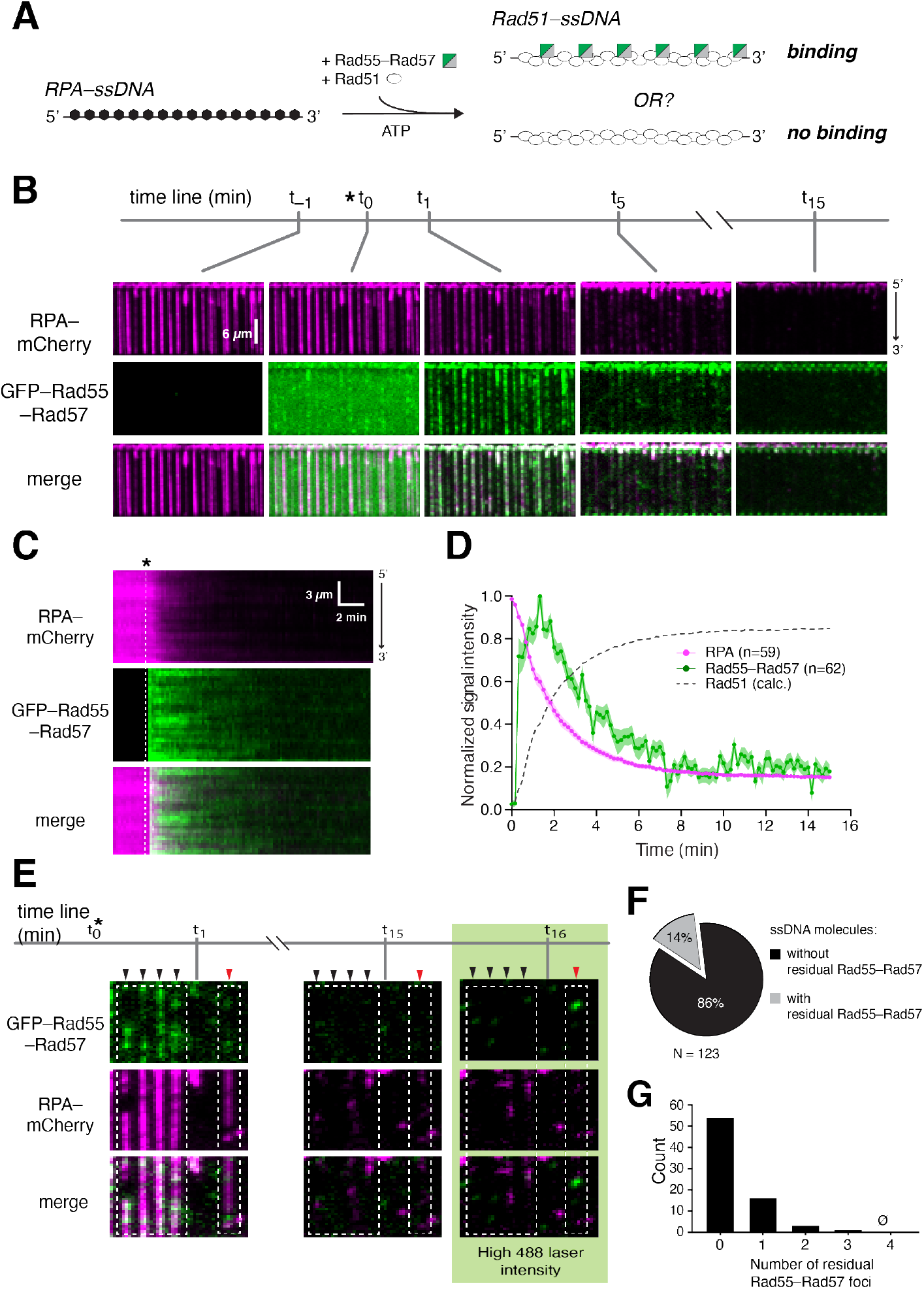
Rad55–Rad57 binds in a transient burst during Rad51 filament assembly. (**A**) Schematic for the experimental setup. (**B**) Wide–field view of RPA–mCherry (magenta) bound ssDNA at indicated time points. Rad51 (unlabeled) and GFP–tagged Rad55–Rad57 (green) injection (t_0_) is highlighted with an asterisk (*). (**C**) Kymograph showing loss of RPA–mCherry after Rad51 and GFP–Rad55–Rad57 injection. (**D**) Mean normalized intensities for RPA–mCherry and GFP–Rad55–Rad57; shaded area represents 95% CI. Calculated Rad51 assembly kinetics is plotted as [1 – RPA] signal for comparison. (**E**) Wide–field view of RPA–mCherry–ssDNA at indicated time points. After 15 min, 488 nM laser power was increased to 200 mW to visualize residual GFP–Rad55–Rad57. Black arrowheads mark ssDNA molecules with no residual Rad55– Rad57, and red arrowhead marks a ssDNA molecule with residual Rad55–Rad57. (**F**) Proportion of ssDNA molecules with residual GFP–Rad55–Rad57 at the 15 min time point. (**G**) Histogram depicting the number of residual GFP–Rad55–57 foci observed per ssDNA molecule at the 15 min time point. See also Figure S2, S3.

To eliminate the possibility that the loss of GFP–Rad55–Rad57 was due to photobleaching, we conducted the experiment at varying laser illumination times. The transient behavior of GFP– Rad55–Rad57 was unchanged, confirming that loss of GFP was not due to photobleaching (Figure S3A). As an additional control, we also quantified the binding kinetics of Hed1–GFP, which is a meiosis–specific protein that binds tightly to Rad51–ssDNA and does not dissociate over the time scales of our measurements (Brown and Bishop, 2014; Busygina et al., 2008; Crickard et al., 2018b). As anticipated, the binding of Hed1–GFP to the Rad51 filaments during their assembly paralleled the dissociation of RPA–mCherry from the ssDNA, but in striking contrast to Rad55– Rad57, Hed1 remained tightly bound to the mature Rad51–ssDNA filaments with no significant loss of the Hed1–GFP signal (Figure S3B). Together, our control experiments confirmed that loss of GFP–Rad55–Rad57 was not due to GFP photobleaching.

Finally, we found that Rad55–Rad57 did not interact with the meiosis–specific recombinase Dmc1 (Brown and Bishop, 2014), indicating that the interactions we have observed are specific for Rad51 (Figure S4). The absence of an interaction between Rad55–Rad57 and Dmc1 is consistent with the inability of Srs2 to strip Dmc1 from ssDNA, suggesting that Dmc1 filaments would have no need for regulation by Rad55–Rad57 (Crickard et al., 2018a; Sasanuma et al., 2013). The results presented above suggest that Rad55–Rad57 binds only transiently to Rad51 filaments during the filament assembly phase and does not form a stable co–filament with Rad51.

### Rad55–Rad57 stimulates Rad51 filament assembly

During HR, Rad51 must displace RPA from ssDNA to form Rad51 filaments, and *in vivo* this process requires the action of mediators such as Rad55–Rad57. We therefore asked how Rad55– Rad57 affects RPA displacement from ssDNA to facilitate Rad51 filament assembly. These assays were conducted at a substoichiometric ratio of Rad55–Rad57 to Rad51 to mimic the ~1:10 physiological ratios of these proteins (Sung, 1997). Strikingly, this sub–stoichiometric amount of Rad55–Rad57 (60 nM) stimulated Rad51 (2 μM) filament assembly ~3–fold, as measured by the loss of RPA–mCherry, yielding a *k*_off,RPA_ of 0.493 ± (0.485 – 0.501; 95% CI) min^−1^ compared to 0.187 ± (0.181 – 0.193; 95% CI) min^−1^ for reactions without Rad55–Rad57 (Figure 3A–C, Table S1). We then measured RPA–Cherry signal remaining on the ssDNA once Rad51 filament assembly had plateaued (15 min) and found that the inclusion of Rad55–Rad57 led to ~2–fold increase in overall RPA displacement by Rad51 (Figure 3D). Our single–molecule results thus show that Rad55–Rad57 strongly stimulates the rate of Rad51 filament assembly at physiological ratios of Rad51 to Rad55–Rad57 and also leads to more extensive coverage of ssDNA by Rad51.

**Figure 3.**
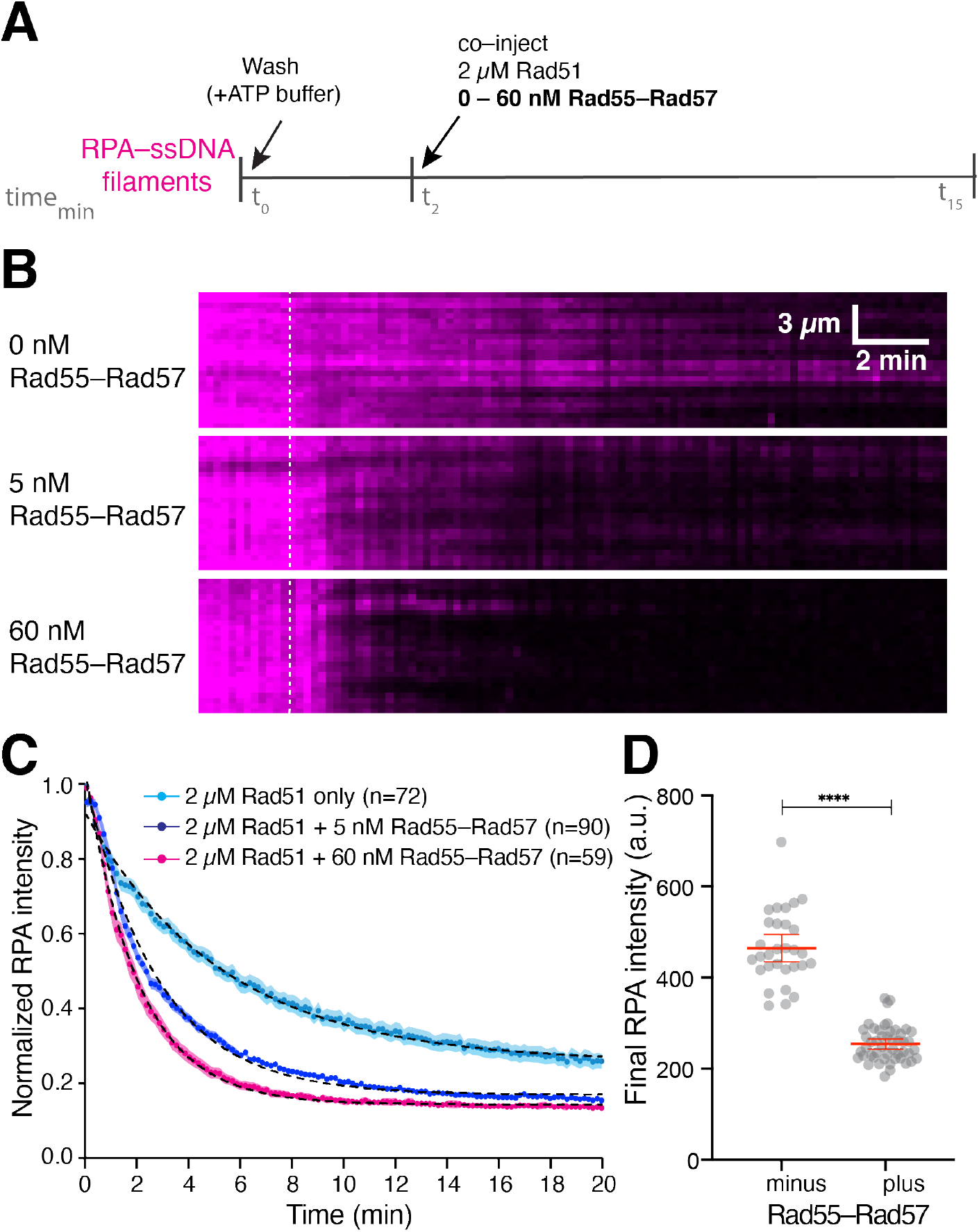
Rad55–Rad57 stimulates Rad51 filament assembly and promotes depletion of RPA. (**A**) Schematic for the experimental setup. (**B**) Kymographs showing loss of RPA–mCherry after Rad51 injection with 0, 5 or 60 nM GFP–Rad55–Rad57. Time of injection is indicated by a white dashed line. (**C**) Kinetics of RPA–mCherry dissociation under indicated conditions. Data is represented as mean normalized RPA intensity; shaded area represents 95% CI. Single exponential decay fits are depicted by a black dotted line. (**D**) RPA–mCherry signal remaining on the ssDNA molecules after Rad51 assembly plateaued, without (N = 30) or with 5 nM Rad55–Rad57 (N = 48). Red bars indicate mean ± 95% CI.

### ATP hydrolysis by Rad55–Rad57 is required for its release from Rad51–ssDNA filaments

Notably, cells expressing the ATP hydrolysis deficient mutants Rad55–K49R or –K49A are sensitized to IR, whereas cells that express the equivalent ATP hydrolysis deficient *rad57* mutants are not, strongly suggesting that the ATP hydrolysis activity of Rad55 is crucial for *in vivo* function of the Rad55–Rad57 complex (Johnson and Symington, 1995). Given that Rad55 shares the conserved Rad51/RecA ATPase core domain of Rad51, we hypothesized that ATP hydrolysis by Rad55 may be required for binding and/or release of Rad55–Rad57 from Rad51–ssDNA filaments during their assembly. We were unable to purify Rad55–Rad57 containing the Rad55–K49R or – K49A mutants due to a lower stability of the corresponding mutant heterodimers. Therefore, as an alternative means to test our hypothesis, Rad51 and Rad55–Rad57 were co–injected with the non– hydrolysable ATP analog AMP–PNP into a sample chamber containing ssDNA molecules bound by RPA–mCherry (Figure 4A). Under these conditions, Rad55–Rad57 associated rapidly with the Rad51 filaments during their assembly and remained irreversibly trapped on the Rad51 filaments throughout the reaction (Figure 4B–D). Notably, Rad55–Rad57 stimulated Rad51–ssDNA filament assembly ~2–fold in reactions with AMP–PNP, even though it remained trapped on the DNA, indicating that Rad55–Rad57 can at least partially stimulate Rad51 filament assembly in the absence of ATP hydrolysis.

**Figure 4.**
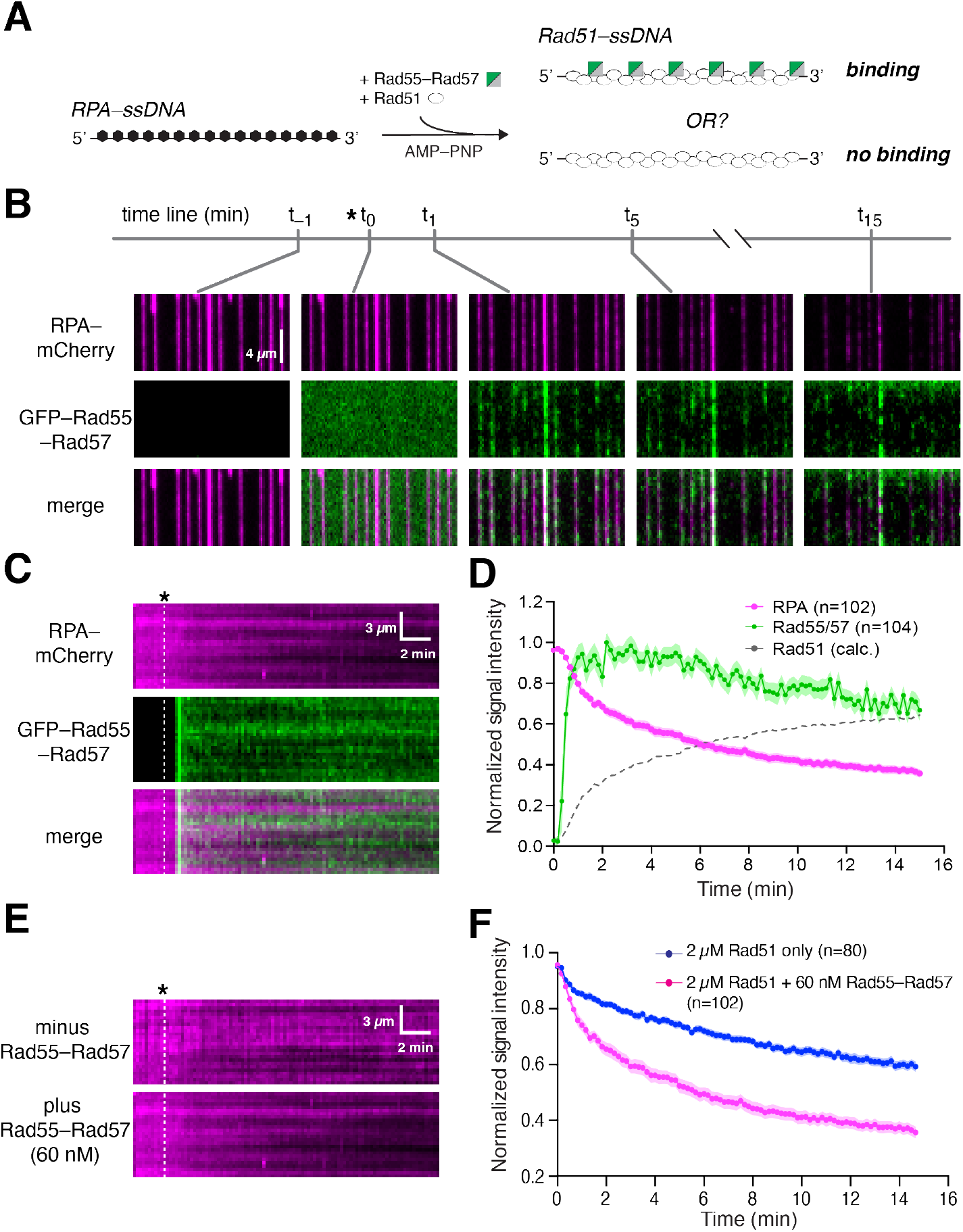
ATP hydrolysis by Rad55–Rad57 is required for its release from Rad51–ssDNA filaments. **(A)** Experimental schematic. (**B)** Wide–field views of RPA–mCherry (magenta) bound ssDNA at the indicated time points. Rad51 and Rad55–Rad57 (green) were injected with 2 mM AMP–PNP. **(C)** Kymograph showing loss of mCherry–RPA and retention of GFP–Rad55–Rad57. **(D)** Mean normalized RPA–mCherry and GFP–Rad55–Rad57 intensity; shaded area represents 95% CI. Rad51 assembly kinetics is plotted as [1 – RPA] signal for comparison. **(E)** Kymographs showing RPA–mCherry loss at the indicated concentrations of Rad55–Rad57 co–injected with 2 μM Rad51. **(F)** Kinetics of RPA–mCherry dissociation under each indicated condition. Data represented as mean normalized intensity; shaded area represents 95% CI. See also Figure S5.

We next tested the ATP hydrolysis deficient mutant Rad51–K191R (Sung and Stratton, 1996) to help determine whether the ATP hydrolysis activity of Rad51 contributes to the turnover of Rad55–Rad57 (Figure S5A). In the presence of 2 mM ATP, Rad55–Rad57 transiently co–localized with Rad51–K191R and then quickly dissociated (*k*_off_ = 0.348 ± (0.305 – 0.393; 95% confidence interval (CI)) min^−1^; Figure S5A–D & Table S1), demonstrating that binding and dissociation of Rad55–Rad57 does not depend on ATP hydrolysis by Rad51. It should be noted that a higher concentration of Rad51–K191R relative to Rad51 (5 μM versus 2 μM, respectively) was required for sufficient filament assembly due to the reduced ssDNA affinity of Rad51–K191R (Fung et al., 2006; Kaniecki et al., 2017), and under these conditions we did not observe further stimulation of Rad51–K191R assembly by Rad55–Rad57 (Figure S5E,F). Together, our results suggest that the ATPase activity of the Rad55–Rad57 complex is required for its release from Rad51 filaments.

### Rad55 ATP hydrolysis regulates turnover of Rad55–Rad57 *in vivo*

To further validate our findings, cells expressing GFP–Rad55, GFP–Rad55–K49R or Rad57– K131R were analyzed by interfoci fluorescence redistribution after photobleaching (iFRAP) to observe the *in vivo* dynamics of GFP–Rad55–Rad57 at IR–induced DNA damage foci (Figure 5). We first tested whether our ATPase deficient mutants complemented the IR defect of *rad55Δ rad57Δ* cells. As expected, expression of Rad57–K131R conferred IR resistance similar to the corresponding wild–type complex, whereas expression of GFP–Rad55–K49R conferred only partial IR resistance in *rad55Δ rad57Δ* cells (Figure 5A) (Johnson and Symington, 1995).

**Figure 5.**
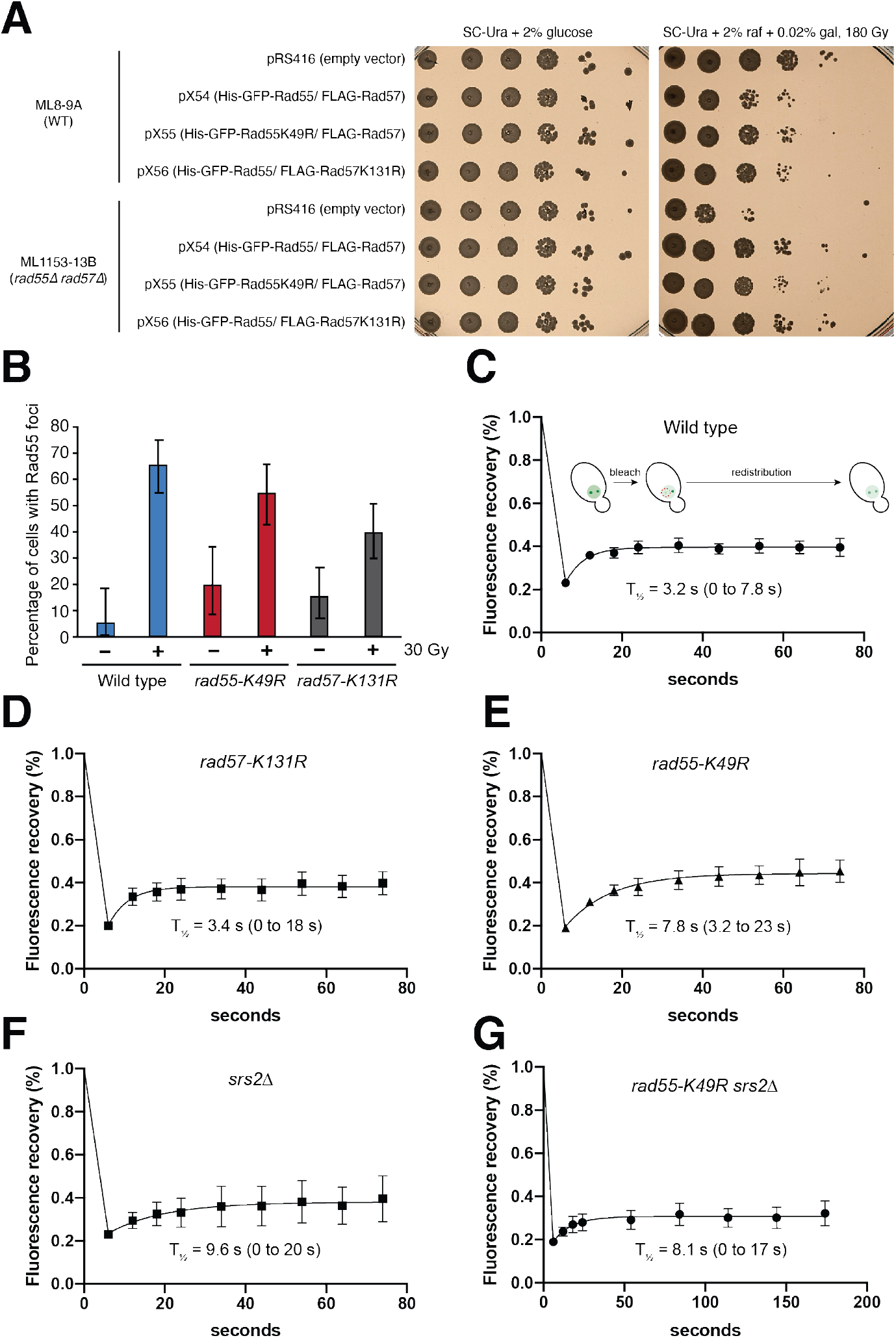
ATP hydrolysis by Rad55 regulates turnover of Rad55–Rad57 *in vivo.* **(A)** WT or *rad55Δrad57Δ* cells were complemented with empty vector (pRS416), or vectors expressing His– GFP–Rad55 and FLAG–Rad57 transgenes as shown (pX54, pX55 and pX56). Ten–fold serial dilutions of indicated strains were plated and subjected to 180 Gy of ionizing radiation (IR). **(B)** Cells were exposed to 30 Gy of IR and inspected for Rad55 foci after 2–hour incubation at 25°C (N = 37–97). *RAD55* and *RAD57* transgenes were expressed from plasmids pX54, pX55 or pX56 in *rad55Δ rad57Δ* cells. These conditions yielded an average of two Rad55 foci per cell. (**C–E)** Cells were exposed to 30 Gy of IR and were subjected to iFRAP, where one Rad55 focus is photobleached and fluorescence redistribution from the other focus monitored by time–lapse microscopy (N = 6–9). (**F–G)** iFRAP was performed as above using a *srs2Δ* deletion strain (strain ML1156–17B).

Our results from the DNA curtain assays predict that Rad55–Rad57 is a highly dynamic component of DNA repair foci. Consistent with this prediction, Rad55–Rad57 turned over rapidly with a half–life of just ~3.2 s, confirming its highly dynamic behavior within DNA repair foci (Figure 5C). The rapid turnover of Rad55–Rad57 in DNA repair foci is in stark contrast to the much slower turnover Rad51, which displays a half of 87 s (Crickard et al., 2018b), providing cellular evidence that Rad55–Rad57 does not form a stable co–filament with Rad51. There was no defect in the turnover of the Rad57–K131R complex, consistent with reported IR resistance of this mutant (Figure 5D) (Johnson and Symington, 1995). In striking contrast, the ATPase deficient GFP–Rad55–K49R mutant complex exhibited a turnover rate that was ~2.4 fold slower than the corresponding wild–type complex, with a half–life of 7.8 s (Figure 5E), suggesting that the ATPase activity of Rad55 helps regulate Rad55–Rad57 dynamics within repair foci. Deletion of *SRS2* also reduced Rad55–Rad57 turnover and a *rad55-K49R srs2Δ* double mutant exhibited similar properties, suggesting that Srs2 may play an active role in removing Rad55–Rad57 from HR intermediates (Figure 5F,G). Together, our data indicate that Rad55–Rad57 is a highly dynamic component of the Rad51–ssDNA presynaptic complex, and similar to Rad51 and Rad52, it is susceptible to removal by Srs2 (De Tullio et al., 2017; Kaniecki et al., 2017).

### Rad55–Rad57 does not block Srs2 translocation

The IR sensitivity of *rad57Δ* cells is partially suppressed by deletion of *SRS2*, and in biochemical assays, Rad55–Rad57 reduces the disruption of Rad51 filaments by Srs2 (Fung et al., 2009; Liu et al., 2011). The current model posits that Rad55–Rad57 associates stably with the Rad51 filament and physically blocks Srs2 translocation, thereby protecting Rad51 filaments from disruption by Srs2. To reconcile our observations with the current model, we first considered the possibility that Rad51 filaments assembled in the presence of Rad55–Rad57 were somehow resistant to Srs2 translocation and disruption. We assembled Rad51–ssDNA filaments either with, or without Rad55–Rad57, flushed out unbound proteins from the sample chamber, and injected mCherry– Srs2. Full length Srs2 is prone to aggregation, so we used the well characterized C–terminal deletion mutant Srs2(1–898) that retains the translocase, Rad51 disruption and ATP hydrolysis activities (Antony et al., 2009; Colavito et al., 2009; Qiu et al., 2013), and we confirmed that it retains its interaction with Rad55–Rad57 (Figure S1). We found that Srs2 was able to translocate on Rad51 filaments formed in the presence of Rad55–Rad57, with only a modest (10.6%) decrease in velocity compared to Rad51 filaments assembled in the absence of Rad55–Rad57 (Figure S6A–C, Table S2). We attribute this reduction in Srs2 velocity to our observation that filament assembly reactions conducted in the presence of Rad55–Rad57 lead to greater coverage of the ssDNA by Rad51 (see Figure 3D).

Next, we asked whether residual Rad55–Rad57 that remained bound on a minority of the Rad51–ssDNA molecules (~14%) after Rad51 filament assembly (Figure 2E–G) could perhaps block Srs2 translocation. We prepared Rad51–ssDNA filaments in the presence of GFP–Rad55– Rad57, flushed out unbound proteins from the sample chamber, and then injected Srs2 in buffer supplemented with RPA–mCherry and 2 mM ATP. As described previously, the 3’→5’ translocation of Srs2 can be visualized by the rapid binding of mCherry–RPA to ssDNA exposed upon Rad51 filament disruption (De Tullio et al., 2018; De Tullio et al., 2017; Kaniecki et al., 2017). Images were acquired using maximum GFP laser illumination (488 nm, 200mW) to clearly visualize any residual GFP–Rad55–Rad57 complexes remaining on the Rad51–ssDNA filaments. As most Rad51–ssDNA filaments did not have any residual Rad55–Rad57 complexes (Figure 2E– G), encounters between translocating Srs2 molecules and Rad55–Rad57 complexes were rare. However, in all cases where Srs2 encountered Rad55–Rad57, it quickly removed or displaced the latter from the Rad51–ssDNA filaments, and in no instance was Srs2 translocation blocked by Rad55–Rad57 (Figure S6D, E). These data demonstrate that residual Rad55–Rad57 complexes bound to mature Rad51 filaments do not physically obstruct Srs2 translocation.

### Competition between RPA and Rad51 for ssDNA binding when Srs2 is present

Given our findings, we considered an alternative model where Rad55–Rad57 acts as a positive regulator of HR by promoting rapid Rad51 filament re–assembly after filament disruption by Srs2 rather than directly blocking Srs2 itself. To test this model, we examined a scenario where RPA and Rad51 must compete with one another for binding to the same ssDNA after the ssDNA is cleared of binding proteins by Srs2 (Figure 6A). In these assays, pre–assembled Rad51–ssDNA filaments were chased with GFP–Srs2 (500 pM) plus RPA–mCherry (500 pM) together with varying concentrations of free Rad51 (0–2 μM; Figure 6A). In the absence of free Rad51, Srs2 displaced the pre–bound Rad51, allowing RPA–mCherry to promptly coat the newly exposed ssDNA (Figure 6B). Previous work has shown that Srs2 is excluded from loading within the interior portions of Rad51 filaments and is instead preferentially loaded onto small clusters of RPA–ssDNA located between adjacent Rad51 filaments (Kaniecki et al., 2017). Consistent with this prior observation, the replacement of Rad51 by RPA allowed for the rapid continuous loading of multiple Srs2 molecules (Figure 6B,C). To quantify this effect, we measured the number of Srs2 molecules translocating through a single point on the ssDNA molecules over a 2.5 min window (Figure S7A, Figure 6C). In the absence of free Rad51, there were an average of 6.6 ± 1.4 Srs2 molecules on each ssDNA (Figure 6B,C). Similar results were obtained in reactions with 500 pM RPA–mCherry plus 0.5 μM free Rad51, indicating that Rad51 was unable to reassemble after the passage of Srs2, even though the free Rad51 concentration was 1,000–fold higher than the free RPA concentration (Figure 6B,C & Table S2). This conclusion was supported by the observation that there was extensive RPA–mCherry rebinding to the ssDNA even though 0.5 μM Rad51 was present in free solution (Figure 6B). In contrast, at 2 μM free Rad51, there was extensive Rad51 filament re–assembly following the initial disruption of the pre–bound Rad51 by Srs2 as evidenced by reduced RPA–mCherry accumulation (Figure 6B), but there was also a 62% decrease in GFP– Srs2 re–loading events (Figure 6B,C). These results are consistent with the genetic observation that the DNA damage sensitivity of *rad55Δ rad57Δ* mutant strains can be suppressed by RAD51 overexpression (Johnson and Symington, 1995), and they also suggest that re–iterative cycles of Rad51 filament re–assembly may progressively suppress Srs2 re–loading by allowing for more extensive Rad51 coverage of the ssDNA (Figure 6B,C).

**Figure 6.**
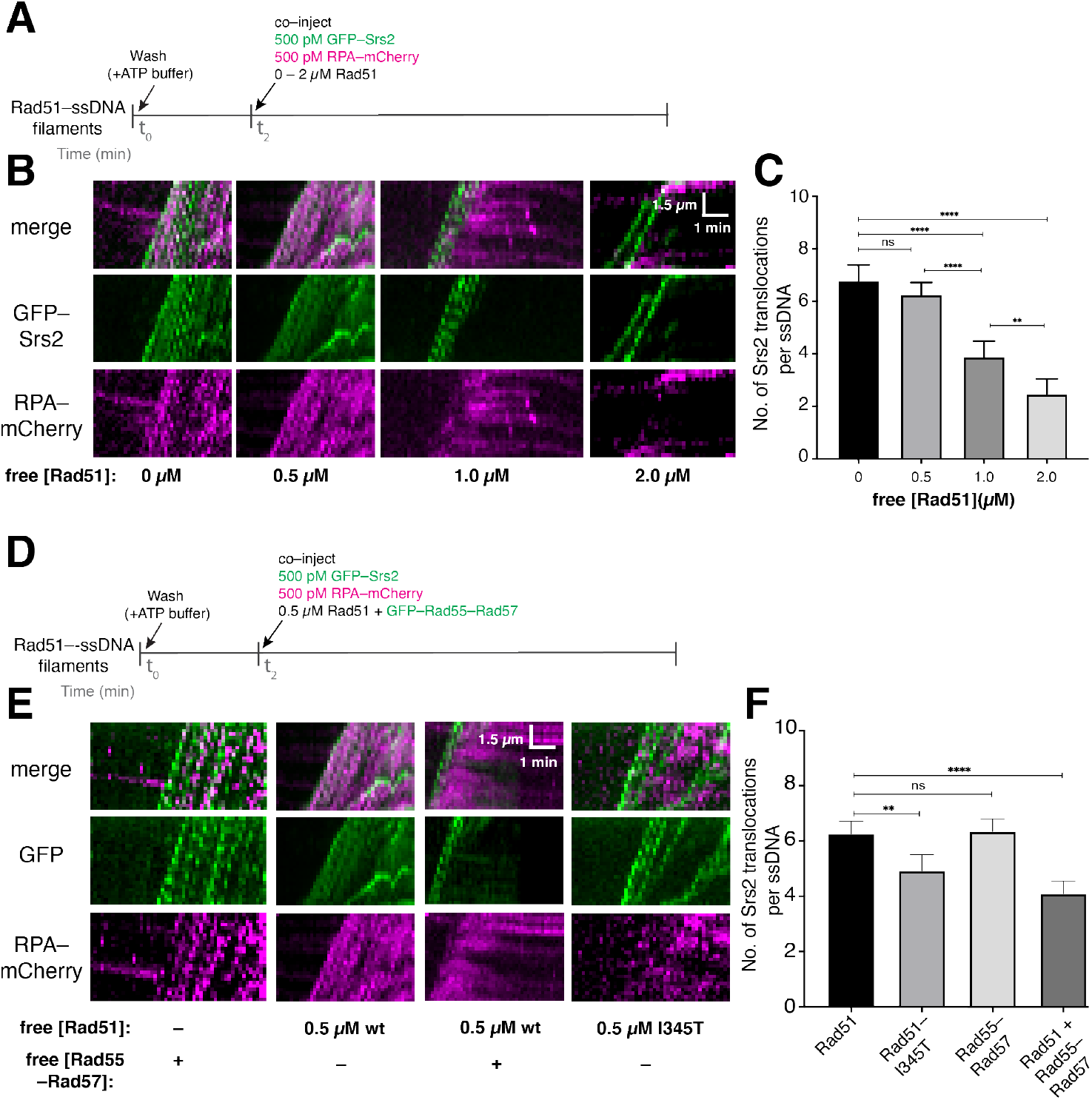
Rad55–Rad57 promotes Rad51 filament re–assembly after disruption by Srs2. (**A**) Assay for assessing competition between Rad51 and RPA in the presence of Srs2. (**B**) Kymographs showing GFP–Srs2 (green) translocation on Rad51–ssDNA and RPA–mCherry (magenta) in the presence of varying concentrations of free Rad51. (**C**) Number of Srs2 translocation events per ssDNA. Error bars indicate 95% CI. (**D**) Assay for assessing competition between Rad51 and RPA in the presence of Rad55–Rad57. (**E**) Kymographs showing GFP–Srs2 and mCherry–RPA re–binding on Rad51–ssDNA with free Rad51 and GFP–Rad55–Rad57; note, 0.5 μM wt Rad51 panels (minus Rad55–Rad57) are reproduced from (B) for comparison. (**F**) Number of Srs2 translocation events per ssDNA under each indicated condition. Error bars indicate 95% CI. See also Figure S7.

### Rad55–Rad57 promotes Rad51 filament reassembly after passage of Srs2

We next asked whether Rad55–Rad57 influences the competition between Rad51 and RPA for ssDNA binding (Figure 6D). For these experiments, we chose a concentration of free Rad51 (0.5 μM) that was insufficient for filament re–assembly when in competition with 500 pM RPA (Figure 6B). Remarkably, inclusion of 60 nM Rad55–Rad57 reduced the amount of RPA–mCherry that accumulated on the ssDNA and also reduced the number of Srs2 binding events by 32% within a 2.5 min window (Figure 6E,F & Table S2). These findings suggest that the presence of Rad55– Rad57 favors Rad51 filament re–assembly in the presence of free RPA after initial disruption by Srs2.

*RAD51–I345T* was isolated as a suppressor mutation that alleviates the sensitivity of *rad55Δ* cells to DNA damage (Fortin and Symington, 2002). Rad51–I345T filaments assemble more rapidly than wild–type Rad51 but are not resistant to Srs2 disruption (Kaniecki et al., 2017). Consistent with our model and previous genetic studies (Fortin and Symington, 2002), Rad51– I345T rapidly re–assembled after disruption by Srs2 in the absence of Rad55–Rad57, leading to a lower RPA–mCherry signal and a corresponding reduction in Srs2 re–loading events (Figure 6E,F & Table S2). Thus, Rad51–I345T overcomes the need for Rad55–Rad57 by allowing for faster Rad51 filament re–assembly. Together, our competition assays demonstrate that Rad55–Rad57 promotes rapid Rad51 filament re–assembly after Srs2 disruption despite the presence of RPA and helps to restrict further Srs2 binding events.

## Discussion

Our work establishes that the *S. cerevisiae* Rad51 paralog complex Rad55–Rad57 interacts with Rad51 only during the earliest stages of Rad51 filament assembly, then promptly dissociates during filament maturation through a mechanism linked to ATP hydrolysis by Rad55. Given the transient nature of this interaction, our work defines Rad55–Rad57 as fulfilling role akin to that of a classical chaperone (Figure 7). The highly transient nature of this interaction may be broadly conserved among eukaryotes, as Bélan *et al.* report similar findings for *C. elegans* RAD51 paralogs in an accompanying paper. Our studies also demonstrate that Rad51 can undergo repeated cycles of assembly and disassembly, the relative rates of which are defined by the opposing actions of Rad55–Rad57 and Srs2 (Figure 7). Continuous remodeling of Rad51 filaments by the combined actions of Rad55–Rad57 and Srs2 could allow for tight quality control during filament assembly and help funnel intermediates through the most appropriate pathway for repair. This model offers a new paradigm for understanding how defects in human RAD51 paralogs and anti–recombinases may influence genome integrity and lead to oncogenic cell transformation.

**Figure 7.**
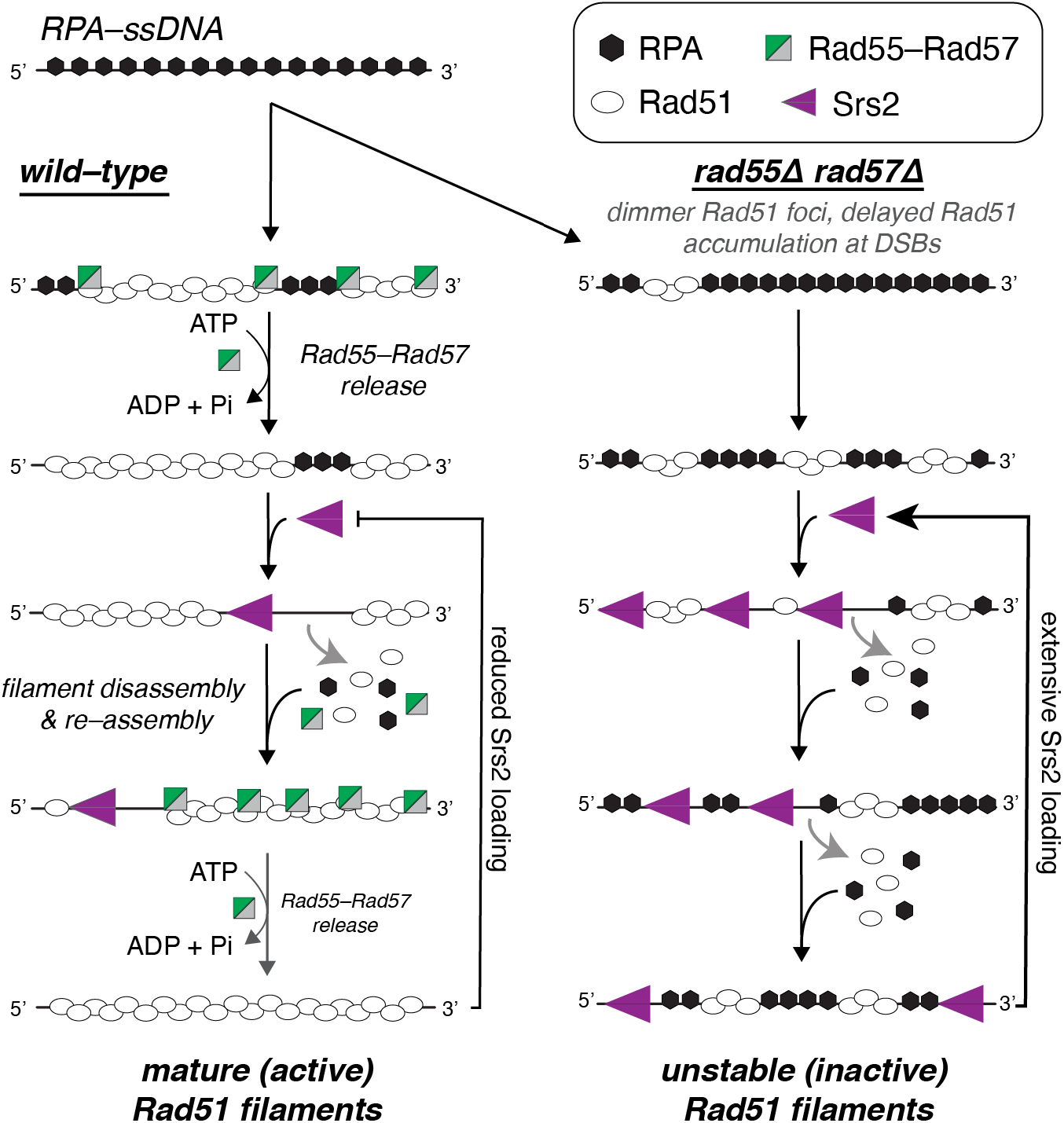
Model for the relationship between Rad55–Rad57 and Srs2. In wild–type cells, Rad55–Rad57 stimulates Rad51 filament assembly through transient binding interactions and then quickly dissociates when Rad55 hydrolyzes ATP. The resulting Rad51 filaments are disrupted by Srs2, which preferentially loads at short tracts of RPA between Rad51 filaments (Kaniecki et al., 2017). New Rad51 filaments are then re–assembled behind Srs2 through the stimulatory action of Rad55–Rad57. Iterative cycles of filament assembly and disassembly may lead to more extensive coverage of the ssDNA by the Rad51 filaments, which in turn helps restrict new Srs2 loading events. In *rad55Δ rad57Δ* cells, filament disruption is favored because RPA can out compete Rad51 for ssDNA binding sites, which also promotes reiterative Srs2 loading resulting in continuous Rad51 filament disruption.

### Antagonistic relationship between Srs2 and Rad55–Rad57

Current knowledge of Rad55–Rad57 posits two independent pro–HR functions for this protein complex. First, as a mediator that relieves RPA inhibition on Rad51 filament assembly, and second, as a unique inhibitor of Srs2 that blocks its translocation via a direct protein–protein physical contact (Gaines et al., 2015; Liu et al., 2011; Sung, 1997). In contrast, our data show that Rad55–Rad57 does not directly inhibit Srs2 translocase activity, but rather counters Srs2 action by promoting rapid Rad51 filament re–assembly (Figure 7). Thus, we demonstrate that the robust mediator activity of Rad55–Rad57 also underlies its role as an antagonist of Srs2. The functional relevance of the protein–protein interaction between Rad55–Rad57 and Srs2 remains unclear, and may affect other aspects of Srs2 function, such as checkpoint activation (Leon Ortiz et al., 2011). However, the amino acid residues at the putative interaction interface between Rad55–Rad57 and Srs2 remain unknown and will have to be identified in order to address its *in vivo* relevance.

Interestingly, there are several parallels between the two major mediator complexes in *S. cerevisiae*, Rad55–Rad57 and Rad52, with respect to their relationship with Srs2. *RAD52* deficient cells are also defective in Rad51 focus formation in response to IR, and this defect can be partially suppressed by the deletion of *SRS2* (Burgess et al., 2009). In Rad51–mediated strand–exchange assays, Rad52 also opposes Srs2 action (Burgess et al., 2009), while being highly susceptible to Srs2 disruption itself (Burgess et al., 2009; De Tullio et al., 2017). Importantly, the Rad52–Rad51 interaction is required for this effect, suggesting that Rad52 stimulation of Rad51 filament re–assembly underlies the Srs2 antagonism. Together, a common theme emerges where activity of Rad51 mediators plays a prominent role in overcoming the anti–recombinase Srs2 by promoting Rad51 filament re–assembly. However, in striking contrast to Rad55–Rad57, Rad52 binds tightly to RPA–ssDNA and also remains tightly bound as part of the mature Rad51–ssDNA presynaptic complex (Gibb et al., 2014). Thus, although Rad52 and Rad55–Rad57 are both recombination mediators, out data suggest that that act through completely distinctive mechanisms.

### Rad51 presynaptic assembly is a fine balance of opposing actions

Stalled forks and DSB substrates can be repaired via one of multiple pathways depending on the cell cycle phase, DNA lesion types, and DNA damage checkpoint activity (Bhat and Cortez, 2018; Mehta and Haber, 2014). Such a scenario would benefit from maintaining DNA substrates primed for multiple pathways, by allowing dynamic modulation by regulators. Our data support such a model where the Rad51 filaments are continuously remodeled through the combined action of recombination mediators and anti–recombinases, which could allow for better quality control of the filament and poise it for repair through the most appropriate pathway. Interestingly, deletion of *SRS2* in *rad52Δ* cells suppresses the defect in Rad51 focus formation, but the Rad51–DNA complexes in these cells are not proficient for repair, indicating that the continuous cycles of filament assembly–disassembly may serve a quality control mechanism for appropriate HR to proceed (Burgess et al., 2009). There are other salient examples of direct competition between factors for determining DNA repair pathway choice, for example, between HR and non– homologous DNA end joining factors at DSB sites (Kass and Jasin, 2010). Our findings support a model where RPA and Rad51 compete for the same ssDNA substrate, and Rad55–Rad57 tilts the balance in favor of Rad51 by continuously stimulating its reassembly, rather than making the filaments resistant to Srs2 disruption. Consistent with this model, induction of Srs2 overexpression after formation of Rad51 foci *in vivo* reduced Rad51 and increased RPA levels within foci, even in the presence of wild–type levels of Rad55–Rad57 (Sasanuma et al., 2013).

How does a mature Rad51 filament remain active for HR if it is constantly disrupted by Srs2? Based on our single molecule results, we propose that rapid reassembly of Rad51 onto ssDNA may help to reduce further Srs2 re–binding. This suppression may arise because more extensive Rad51 assembly leads to fewer RPA–bound gaps within the Rad51 filaments. This may help create a conducive environment for Rad51 filament assembly and maintenance because RPA–bound regions represent preferred loading sites for Srs2 (Kaniecki et al., 2017). Consistent with the idea that Srs2 is recruited to RPA–ssDNA is the finding that Srs2 is recruited to recombination foci even in the absence of Rad51 (Burgess et al., 2009). It is also possible, that by stimulating Rad51 assembly sufficiently, Rad55–Rad57 allows a critical level of Rad51–ssDNA to accumulate *in vivo*, enabling downstream Rad51–dependent factors to bind these presynaptic filaments, which in turn may inhibit or exclude Srs2.

### What intermediate does Rad55–Rad57 act upon?

Our data show that Rad55–Rad57 binds to Rad51 filaments only during the earliest stages of their assembly and then quickly dissociates through an ATP hydrolysis–dependent mechanism as the filaments mature (Bianco et al., 1998; Kowalczykowski, 2015). Interestingly, in our ssDNA curtain assays, a ratio of 60 nM Rad55–Rad57 to 2 μM Rad51 (mimicking the native ~1:10 ratio of these proteins *in vivo (Sung, 1997))* yielded a 3–fold enhancement in filament assembly, whereas a 12–fold increase in Rad55–Rad57 concentration (from 5 nM to 60 nM) did not yield a concomitant increase in filament assembly rates even though free Rad51 was present in a large molar excess (~30–fold excess) (Figure 3C). This result suggests that Rad55–Rad57 primarily, or exclusively, acts on a Rad51–ssDNA intermediate, which would be limiting in our assays. Consistent with this hypothesis, Rad51 does not co–immunoprecipitate with Rad55–Rad57 from yeast cell extracts, and Rad51 interacts with Rad55–Rad57 only weakly in pull–down assays (Liu et al., 2011; Sung, 1997). In addition, *RAD55* and *RAD57* deletion mutants show a cold sensitive phenotype (Lovett and Mortimer, 1987), unique among the RAD52 epistasis group members, leading to the suggestion that Rad55–Rad57 enhances the stability of an otherwise unstable intermediate (Johnson and Symington, 1995). What is the nature of the intermediate targeted by Rad55–Rad57? Rad51 filament assembly proceeds via a rate–limiting nucleation step where short Rad51 clusters bind ssDNA in an unstable manner, resulting in many non–productive nucleation events. A fraction of these nucleation events allow for elongation by Rad51 subunit addition to the filament ends (Kowalczykowski, 2015). Our results suggest that the target of Rad55–Rad57 is enriched early during filament assembly and depleted at later time points. Considering that Rad51 paralogs have been proposed to bind Rad51–ssDNA filament ends (Liu et al., 2011; Taylor et al., 2016), one can imagine a model where Rad55–Rad57 binds the unstable Rad51 nucleation clusters on ssDNA, thereby stabilizing this intermediate and increasing the overall rate of Rad51 filament assembly. As the filament assembly reaction proceeds, there is a concomitant reduction in the density of Rad51–ssDNA filament ends, leading to the loss of Rad55–Rad57 binding targets as the filaments mature over time.

### Rad51 paralogs as molecular chaperones

Using real–time single–molecule imaging we have demonstrated that the Rad55–Rad57 complex associates with Rad51 filaments during a transient burst that coincides with the earliest stages of filament assembly and then rapidly dissociates during filament maturation. Companion cell biological analysis has verified that Rad55–Rad57 behaves as a highly dynamic component of DSB repair foci *in vivo*. We propose therefore that Rad55–Rad57 acts as a classical molecular chaperone to help promote Rad51 filament nucleation, and our findings lead to a new framework for understanding the behavior and function of Rad51 paralogs in HR (Figure 7). This chaperone–like behavior of the Rad51 paralogs is likely broadly conserved among eukaryotes, as Bélan *et al.* report very similar observations with the *C. elegans* Rad51 paralogs. Given the importance of Rad51 paralogs in human health and disease, it would be importantly to determine whether the human Rad51 paralog complexes also act as molecular chaperones, and how cancer mutations alter this behavior and function.

## STAR ★ METHODS

Detailed methods are provided in the online version of this paper and include the following:

- KEY RESOURCES TABLE
- LEAD CONTACT AND MATERIALS AVAILABILITY
- METHOD DETAILS

- Purification of GFP–Rad55–Rad57
- Purification of Rad51, RPA, Srs2, Hed1 and Dmc1
- Camptothecin sensitivity assays
- IR sensitivity assays
- Live cell imaging of Rad55–Rad57 foci
- Affinity pull down assays
- Single molecule DNA curtain assays
- Rad51 filament assembly and kinetics
- In vitro photobleaching controls and Hed1–binding assays
- Srs2 antirecombinase assays
- Single–molecule data collection and analysis
- Rad51 assembly kinetics
- Analysis of residual Rad55–Rad57
- Analysis of Srs2 trajectories
- Statistical information
- DATA AND CODE AVAIBILITY

## SUPPLEMENTAL INFORMATION

Supplemental Information includes 7 Figures and 2 Tables.

## Acknowledgments

We thank Rodney Rothstein and members of the Greene and Sung laboratories for discussion and comments on the manuscript. We thank Simon Boulton, David Rueda, Ondrej Belan and colleagues for sharing data prior to publication and for comments on our manuscript. This research was funded by NIH grants R35GM118026 (E.C.G.), R35GM126997 (L.S.), R35CA241801, RO1ES007061 (P.S.), PO1CA092584 (P.S. and E.C.G.), a Gray Foundation Team Science Grant under the Basser Initiative (P.S.), and a Wellcome Trust Collaborative Award in Science (Grant. No. 206292/D/17/Z; E.C.G.). M.L. was supported by the Danish Council for Independent Research, the Villum Foundation, and the Danish National Research Foundation (DNRF115). P.S. is the recipient of a CPRIT REI Award (RR180029) and holder of the Robert A. Welch Distinguished Chair in Biochemistry (AQ–0012).

## Author Contributions

U.R. cloned and purified GFP–tagged Rad55–Rad57, designed and conducted all single molecule assays and conducted bulk biochemical assays. Y.K. assisted with Rad55–Rad57 expression and purification. L.M. conducted genetic complementation experiments. M.L. conducted all *in vivo* measurements of Rad55–Rad57 foci. U.R. and E.C.G. co–wrote the manuscript with input from all other co–authors.

## STAR ★ METHODS

### RESOURCE AVAILABILITY

#### Lead Contact

Further information and requests for resources and reagents should be directed to and will be fulfilled by the Lead Contact, Eric C. Greene.

#### Materials Availability

All reagents generated in this study are available from the Lead Contact with a completed Materials Transfer Agreement.

#### Data and Code Availability

All kymographs used as the original source for data analysis throughout the paper are available on Mendeley *[DOI link will be provided upon acceptance].*

### EXPERIMENTAL MODEL AND SUBJECT DETAILS

For single molecule and bulk biochemical assays *S. cerevisiae* His_6_–GFP–Rad55 and FLAG– Rad57 were co–expressed in *S. cerevisiae* strain yRP654 *(MATa ura3–52 trp1* Δ *leu2Δ1 his3Δ2OO pep4::HIS3 prb1Δ1.6R can1 GAL;* gift from the L. Prakash laboratory (Wilson et al., 2013)) in synthetic media lacking uracil (Sherman, 1991), 2% galactose, 3% lactic acid, 3% glycerol at 30°C for 30 hr. *S. cerevisiae* Rad51, Dmc1, RPA, RPA–mCherry, GFP–Srs2, mCherry–Srs2, Hed1– GFP were all overexpressed in *E. coli* BL21 (DE3) Rosetta2 cells (EMD Millipore Cat# 714003; F– *ompT hsdS*B(rB– mB–) *gal dcm* (DE3) pRARE2 (CamR)) grown in Luria Broth. Growth temperature, expression and purification information for each different protein are provided below in the METHOD DETAILS.

Camptothecin sensitivity assays used the isogenic strains, LSY0697 *(MATa met17–s, leu2–3,112, trp1–1, ura3–1, can1, his3-11,15*) and LSY0848 *(MATa rad55::LEU2 rad57::LEU2 met17–s leu2–3,112 trp1–1 ura3–1 can1 his3-11,15*), were transformed with the pESC–URA encoding His_6_–GFP–RAD55 and FLAG–RAD57 and containing the *URA3* marker. Ura+ transformants were selected on synthetic complete plates lacking uracil (SC–Ura) + 2% glucose. Additional details are provided below in the METHODS DETAILS.

For live cell imaging, GFP–tagged Rad55, Rad57 and mutants thereof were expressed from the bi–directional *GAL1–10* promoter on plasmids pESC–URA3–His–GFP– Rad55/FLAG–Rad57, pESC–URA3–His–GFP–Rad55–K49R/FLAG–Rad57 or pESC–URA3–His–GFP– Rad55/FLAG–Rad57–K131R in *rad55Δ rad57Δ* (ML1153–13B; *MAT*a *rad55Δ rad57::LEU2)* or *rad55Δ rad57Δ srs2Δ* (ML1156–17B; *MATa rad55Δ rad57::LEU2 srs2::HIS3)* budding yeast strains cultured at 25°C in synthetic complete medium lacking uracil (SC–Ura) containing 2% raffinose as a carbon source and 0.02% galactose. Additional details are provided below in the METHODS DETAILS.

Plasmids for cloning were grown in *E. coli* DH5alpha (NEB Cat# C2987H; *fhuA2 a(argF–lacZ) U169 phoA glnV44 a80a(lacZ)M15 gyrA96 recA1 relA1 endA1 thi–1 hsdR17)* or *E. coli* NEB Turbo (NEB Cat# C2984H; *FproA+B+ lacIqD lacZM15/fhuA2 D(lac–proAB) glnVgal R(zgb 210::Tn10)TetSendA1 thi–1 D(hsdS–mcrB)5*).

### METHOD DETAILS

#### Purification of GFP–Rad55–Rad57

A GFP tag was cloned into pESC–URA vector containing (His)_6_–tagged RAD55 and FLAG– tagged RAD57 (Gaines et al., 2015), to generate His_6_–GFP–RAD55 and FLAG–RAD57. Cloning was done by the InFusion HD cloning system (Takara Biotech, Cat No. 638920) using the manufacturer’s instructions. *S. cerevisiae* expressing His_6_–GFP–Rad55–FLAG–Rad57 were collected by centrifugation to obtain an 85 g cell pellet. All steps were carried out at 4°C, and 2 mM ATP and 5 mM Mg^2+^ was maintained in all buffers during lysis and purification steps to minimize the aggregation and precipitation of GFP–Rad55–Rad57. The cell pellet was lysed using a bead beater in 250 ml of CBB buffer (50 mM Tris–HCl [pH 8.0], 600 mM KCl, 10% sucrose, 2 mM EDTA, 0.01% IGEPAL CA–630 (Sigma, Cat No. I8896), 1 mM DTT, 1 mM phenylmethylsulfonyl fluoride, and 5 μg ml^−1^ each of protease inhibitors aprotinin, chymostatin, leupeptin and pepstatin), supplemented with 2 mM ATP and 5 mM MgCl2. Lysate was clarified by ultracentrifugation and proteins precipitated with 40% ammonium sulphate. The protein pellet was resuspended in 30 ml of buffer K (20 mM K_2_HPO_4_ [pH 7.5], 10% glycerol, 0.5 mM EDTA, 1 mM DTT, 0.01% IGEPAL, 2 mM ATP, 5 mM MgCl_2_, 5 μg ml^−1^ each of protease inhibitors) supplemented with 300 mM KCl and 20 mM imidazole, and incubated with 2–ml Ni–NTA affinity resin (Qiagen, Cat No. 30210) for 1 hr with gentle mixing. The resin was washed with 50 ml of buffer K with 1000 mM KCl and 30 ml of buffer K with 300 mM KCl, followed by protein elution with a total 20 ml of buffer K supplemented with 300 mM KCl and 200 mM imidazole. Fractions containing GFP–Rad55–Rad57 were pooled and incubated with 1.5–ml anti–FLAG affinity resin (Sigma, Cat No. A2220) for 1 h, with gentle mixing. The resin was washed with 50 ml of buffer K with 1000 mM KCl and 30 ml of buffer K with 300 mM KCl, followed by protein elution with a total of 9 ml of buffer K supplemented with 300 mM KCl and 0.2 mg ml^−1^ FLAG peptide (Sigma). GFP–Rad55–Rad57 containing fractions were pooled and further purified by gel filtration using a 24 ml Superdex 200 column (GE Healthcare, Cat No. 29321906) in buffer K supplemented with 300 mM KCl. The peak fractions of GFP–Rad55–Rad57 were pooled, concentrated in an Amicon Ultra micro–concentrator (Millipore, Cat No.: UFC803024), snap– frozen in liquid nitrogen and stored at –80°C. The yield of purified GFP–Rad55–Rad57 was 900 μg.

#### Purification of Rad51, RPA, Srs2, Hed1 and Dmc1

Rad51, RPA, RPA–mCherry, GFP–Srs2, mCherry–Srs2, Hed1–GFP and Dmc1 were purified as previously described (Crickard et al., 2018a; Crickard et al., 2018b; Crickard et al., 2020; De Tullio et al., 2017; Kaniecki et al., 2017). In brief, 6xHis–SUMO–Rad51 was overexpressed in *E. coli* BL21 (DE3) Rosetta2 cells at 37°C to an OD600 of 0.4–0.6. Expression was induced by addition of 0.5 mM IPTG for 3 hours at 37°C. Overexpression cells were harvested and stored at –80°C. Cells were lysed by freeze–thaw in Cell Lysis Buffer (30 mM Tris–HCl [pH 8.0], 1 M NaCl, 10% glycerol, 10 mM imidazole, 5 mM BME and protease inhibitor cocktail (Roche, Cat. No. 05892953001)). Crude lysates were sonicated for 6 pulses of 30 seconds (s) on and 2 minutes (min) off, and then clarified by ultracentrifugation at 100,000 x g for 45 minutes. The extract was then bound to 1 mL of pre–equilibrated Ni–NTA resin for 1 hour with rotation. The resin was then washed 3X with CLB and eluted in CLB + 200 mM imidazole. The protein was mixed with 400 units of the SUMO protease Ulp1 (Sigma–Aldrich, Cat. No. SAE0067–2500UN) and dialyzed overnight at 4°C into Rad51 buffer (30 mM Tris–HCl [pH 8.0], 150 mM NaCl, 1 mM EDTA, 10% Glycerol, 10 mM imidazole). The 6xHis–SUMO tag and SUMO protease were removed by passing the dialyzed proteins over a second 1 mL Ni–NTA column. The purified Rad51 was then stored at –80°C in single use aliquots.

RPA and RPA–mCherry were overexpressed in *E. coli* BL21 (DE3) Rosetta2 grown at 37°C to an OD600 of 0.4–0.6. The temperature of the cultures was adjusted to 18°C and expression was induced with addition of 0.2 mM IPTG for 8–12 hours. After overexpression cells were harvested and stored at –80°C. Cells were lysed by freeze–thaw in Lysis buffer (40 mM NaHPO_4_ [pH 7.5], 600 mM KCl, 5% glycerol, 5 mM imidazole, 0.1 mM tris–(2–carboxyethyl)phosphine (TCEP), 0.05% Tween 20, and protease inhibitor cocktail [Roche, Cat. No. 05892953001]. Crude lysates were sonicated on ice 6 x 30 s on and 2 min off. The lysate was clarified by ultracentrifugation at 100,000 x g for 45 min. The lysate was bound to pre–equilibrated Talon (Clontech, Cat No. 635503) resin for 30 min at 4°C. The bound protein was washed with nickel A buffer (40 mM NaHPO_4_ [pH 7.5], 300 mM KCl, 5% Glycerol, 5 mM imidazole, 0.02% Tween–20). The protein was then eluted in nickel A buffer + 200 mM imidazole. Peak fractions were then further fractionated on a Superdex 200 size exclusion column equilibrated with SEC buffer (40 mM NaHPO_4_ [pH 7.5], 200 mM KCl, 10% Glycerol, 0.02% Tween–20). Peak fractions, containing all three subunits of RPA, were then pooled and stored in single use aliquots at –80°C. For unlabeled RPA an additional purification step was included in between the Talon column and the size inclusion column. In this case Peak fractions from the Talon column were bound to Chitin resin (NEB, Cat No. S6651L) and then washed in nickel buffer A. The bound protein was then incubated overnight in nickel A buffer plus 100 mM DTT this removed the chitin tag and eluted the RPA protein. This step was followed by the size exclusion step listed above.

For Srs2 expression, a pET11c vector encoding 9xHis–tagged Srs2 and pET15b vectors encoding GFP–Srs2(1–898), mCherry–Srs2(1–898) were introduced into *E*. *coli* Rosetta2 (DE3) cells (Novagen, Cat No. 69450). Cells were grown in 3 L of Luria broth (LB) at 37°C to an optical density 600 (OD600) of 1–2. The temperature was reduced to 16°C before addition of 0.1 mM isopropyl–b–D–thiogalactopyranoside (IPTG). After 20 hr of growth, cells were pelleted and frozen at –80°C. The pellet was then resuspended in lysis buffer (40 mM NaHPO_4_ [pH 7.5], 600 mM KCl, 5% glycerol, 10 mM imidazole [pH 7.8], 0.1 mM tris(2–carboxyethyl)phosphine (TCEP), 0.05% Tween–20, 10 mM E–64, 1 pill per 100 mL of protease inhibitor cocktail tablets (Sigma–Aldrich, Cat. No. C4706–10G), 1 mM benzamidine, 1 mM PMSF, and 0.125% myo– inositol) and lysed by sonication on ice. The lysate was clarified by ultracentrifugation and incubated for 30 min with a Ni–NTA resin (Clontech) that was equilibrated with buffer nickel A (40 mM NaHPO4 [pH 7.5], 300 mM KCl, 5% glycerol, 15 mM imidazole, 0.02% Tween–20, 1 mM benzamidine, 1 mM PMSF, and 0.125% myo–inositol). Before elution, the Ni–NTA column was washed with buffer nickel A. The proteins were eluted with a step of buffer nickel A containing 400 mM imidazole (pH 7.8). The eluate was then dialyzed against heparin buffer (20 mM NaHPO4 [pH 7.5], 100 mM KCl, 5% glycerol, 0.01% Tween– 20, 1 mM TCEP, 2 mM EDTA, and 0.125% myo–inositol) during 2 hr. The eluate was then loaded onto a 5 mL HiTrap Heparin column (GE Lifesciences, Cat No. GE17–0407–01) equilibrated with heparin buffer, and proteins were eluted with a step of heparin buffer containing 500 mM KCl. The purified fraction was applied to a Superdex 200 size exclusion column equilibrated with storage buffer (40 mM NaHPO4 [pH 7.5], 300 mM KCl, 10% glycerol, 0.01% Tween–20, 1 mM TCEP, 0.5 mM EDTA, and 0.125% myo–inositol). Fractions corresponding to monomeric Srs2 were pooled, concentrated, flash frozen in liquid nitrogen, and stored at –80°C.

For Dmc1 expression, the pNRB150 vector harboring N–terminally (His)_6_–tagged ScDmc1 was introduced into BL21[DE3] Rosetta cells (Novagen, Cat No. 69450). An overnight bacterial culture was diluted 50–fold in 2× LB media supplemented with ampicillin (100 μg/ml) and chloramphenicol (34 μg/ml) and grown at 37°C to OD_600_ = 0.8. ScDmc1 expression was induced with 0.1 mM IPTG for 16 h at 16°C. Cell lysate preparation and all the protein purification steps were conducted at 4 °C in buffer T (25 mM Tris–HCl, pH 7.4, 10% glycerol, 0.5 mM EDTA, 0.01% IGEPAL CA–630 (Sigma, Cat No. 9002–93–1), 1 mM DTT) supplemented with 2 mM ATP and 2 mM MgCl2. Chromatographic column fractions were screened for their ScDmc1 content by 12% SDS–PAGE and Coomassie Blue staining. The *E. coli* cell paste (20 g) was suspended in 100 ml of buffer supplemented with 500 mM KCl, 1 mM phenylmethylsulfonyl fluoride, 0.5 mM benzamidine and 5 μg/ml each of aprotinin, chymostatin, leupeptin, and pepstatin. Cells were disrupted by sonication. After ultracentrifugation (100,000 × *g* for 90 min), the lysate was incubated with 2 ml of Talon affinity resin (Clontech, Cat No. 635503) for 2 h with gentle mixing. The matrix was poured into a column with an internal diameter of 1 cm and washed sequentially with 20 ml of buffer with 500 mM KCl and with 150 mM KCl, respectively, followed by ScDmc1 elution using buffer supplemented with 150 mM KCl and 200 mM imidazole. The protein pool was diluted with an equal volume of buffer T and fractionated in a 1 ml Heparin Sepharose column (GE Healthcare, Cat No. 17–0406–01) with a 30 ml gradient of 150–1000 mM KCl, collecting 1 ml fractions. Fractions containing ScDmc1 (eluting at ~500 mM KCl) were pooled, diluted to the conductivity of 150 mM KCl and further fractionated in a 1 ml Mono Q column with a 30 ml gradient of 150–500 mM KCl, collecting 1 ml fractions. Fractions containing ScDmc1 (eluting at ~300 mM KCl) were pooled, concentrated in an Amicon Ultra micro–concentrator (Millipore, Cat No. UFC501096), snap–frozen in liquid nitrogen, and stored at −80°C.

For Hed1 expression, a pGEX plasmid encoding GST–Hed1–6xHis–GFP was transformed into *Escherichia coli* Rosetta (DE3) cells (Novagen, Cat. No. 69450). Cells were grown to an OD of 0.6–0.8 at 37°C, and cultures were then shifted to 16°C and induced overnight with 0.1 mM IPTG. After overnight expression, cells were harvested and re–suspended in 20 ml/l cell lysis buffer (50 mM Tris–Cl [pH 7.5], 700 mM KCl, 1 mM EDTA, 10 mM BME, protease inhibitor cocktail (Roche, Cat. No. 05892988001), 10% glycerol, and 1 mM PMSF). Cells were lysed with lysozyme and sonicated. The lysate was clarified by ultracentrifugation for 45 min at 142,000 x g. Clarified extract was incubated in batch with glutathione resin (GE Healthcare, Cat. No. 17–0756– 01) for 1 h at 4°C. After 1 h, the supernatant was removed and the resin washed with 2 × 10 column volumes (CV) with buffer K1000 (20 mM Tris–Cl [pH 7.5], 1 M KCl, 10 mM BME, 1 mM PMSF, 10% glycerol, 2.5 mM imidazole). Resin was then washed with buffer K300 (20 mM Tris–Cl [pH 7.5], 0.3 M KCl, 10 mM BME, 1 mM PMSF, 10% glycerol, 2.5 mM imidazole). Protein was eluted with buffer K300 plus 25 mM glutathione. Peak fractions were bound to cOmplete Nickel Resin (Roche, Cat. No. 05893682001) for 1 h at 4°C. Resin was then washed 2 × 5 CV of buffer K1000, followed washing with 2 × 5 CV of buffer K300. Protein was then eluted with buffer K300 plus 100 mM imidazole. Peak fractions were pooled and dialyzed against buffer K 150 (20 mM Tris– Cl [pH 7.5], 150 mM KCl, 10 mM BME, 1 mM PMSF, 10% glycerol). Proteins were quantified by absorbance at 280 nm, and in the case of GFP and mCherry, protein concentrations were quantified by measuring the absorbance of the chromophores at 488 nM (e_488 nm_ = 55,000/cm/M) or 587 nm (e_587 nm_ = 72,000/cm/ M), respectively. Samples were flash–frozen and stored at –80°C.

#### Camptothecin sensitivity assays

Isogenic strains, LSY0697 *(MATa met17–s, leu2–3,112, trp1–1, ura3–1, can1, his3–11,15)* and LSY0848 *(MATa rad55::LEU2 rad57::LEU2 met17–s leu2–3,112 trp1–1 ura3–1 can1 his3–11,15*), were transformed with the pESC–URA encoding His6–GFP–RAD55 and FLAG–RAD57 (as described above) and containing the *URA3* marker. Ura+ transformants were selected on synthetic complete plates lacking uracil (SC–Ura) + 2% glucose. Two independent colonies were picked from each transformation and used to inoculate 5 mL SC–Ura + 2% glucose. When cultures reached late log phase, 0.25 mL were pelleted from each culture, washed twice with 1 mL of water and the pellets were re–suspended in 10 mL SC–Ura +2% glucose or 10 mL SC–Ura +1% galactose. Cultures were then incubated at 30°C until they reach OD600 = 0.4 (~5 h in glucose– containing medium and ~20 h in galactose–containing medium). For the cells grown with glucose, 5 μL from 10–fold serial dilutions of each culture were spotted on SC–Ura + 2% glucose plates containing 0, 0.5, 1 or 5 μg/mL camptothecin (CPT). For the galactose cultures, 5 μL of each 10– fold serial dilution was spotted on SC–Ura + 1% galactose plates containing 0, 0.5, 1 or 5 μg/mL CPT. Plates were incubated at room temperature (~23°C) for 2–3 days before imaging.

#### IR sensitivity assays

Ten–fold serial dilutions of overnight cultures of wild–type (ML8–9A) and *rad55Δ rad57Δ* null mutant cells (ML1153–13B) transformed with empty vector (pRS416) or vectors expressing His– GFP–Rad55 and FLAG–Rad57 transgenes as indicated (pX54, pX55 and pX56) were plated on SC–Ura containing either 2% glucose (repression) or 2% raffinose supplemented with 0.02% galactose (induction). The inducing plates were X–irradiated with 180 Gy.

#### Live cell imaging of Rad55–Rad57 foci

Medium for yeast cell culture was prepared as described (Eckert-Boulet et al., 2011). GFP–tagged Rad55, Rad57 and mutants thereof were expressed from the bi–directional *GAL1–10* promoter on plasmids pESC–URA3–His–GFP– Rad55/FLAG–Rad57, pESC–URA3–His–GFP–Rad55– K49R/FLAG–Rad57 or pESC–URA3–His–GFP– Rad55/FLAG–Rad57–K131R in *rad55Δ rad57Δ* (ML1153–13B; *MAT*a *rad55Δ rad57::LEU2*) or *rad55Δ rad57Δ srs2Δ* (ML1156–17B; *MATa rad55Δ rad57::LEU2 srs2::HIS3*) budding yeast strains cultured in synthetic complete medium lacking uracil (SC–Ura) containing 2% raffinose as a carbon source and 0.02% galactose to induce expression of Rad55–Rad57. The strains were isogenic *ADE2 RAD5* derivatives of W303 (Thomas and Rothstein, 1989). Prior to microscopy, transformed strains were grown overnight in SC–Ura containing 2% raffinose and 0.02% galactose with shaking at 25°C, diluted to OD600 = 0.2 and grown for another 3 hours in SC– Ura containing 2% raffinose and 0.02% galactose with shaking at 25°C. Rad55–Rad57 foci were induced by 40 Gy using an X–ray irradiator (Faxitron, Cat. No. CP–160). Interfoci fluorescence redistribution after photobleaching (iFRAP) was conducted as described previously (Klein et al., 2019). In brief, exponentially growing cells were exposed to 40 Gy, which induces an average of 2 DSBs per cell (Lisby et al., 2001), and cultured shaking for another 3 hours at 25°C prior to iFRAP. Cells with two roughly equal size Rad55 foci were selected and one focus photobleached at 488 nm for 0.7 sec at 10% laser intensity. One fluorescence image was acquired before photobleaching and a time series of 9 images after photobleaching using a wide–field microscope (DeltaVision Elite; Applied Precision) equipped with a 100× objective lens with a numerical aperture of 1.35 (U–PLAN S–APO, NA 1.4; Olympus), a cooled EMCCD camera (Evolve 512; Photometrics), and a solid–state illumination source (Insight; Applied Precision, Inc). At each time point, 5 optical sections separated by 0.4 μm were acquired with softWoRx (Applied Precision) software. Processing and quantitative measurements of fluorescence intensities were performed with Volocity software (PerkinElmer).

#### Affinity pull down assays

Purified His_6_–GFP–Rad55 and FLAG–Rad57 complex (150 nM) was incubated with 150 nM His6–Srs2 in buffer P (25 mM Tris–HCl [pH 7.5], 10 mM MgCl2, 50 mM NaCl, 2.5 mM ATP, 1 mM DTT, 10% glycerol and 0.05% NP–40) for 1 h at 25°C. Equilibrated anti–FLAG beads (Sigma, Cat. No. A2220) were added to the mixture and incubated for 1 h. The beads and supernatant were separated by centrifugation, and the beads washed twice with binding buffer P. The protein complexes were eluted by boiling at 95°C for 5 min in SDS–PAGE loading buffer, resolved by a 10% SDS–PAGE gel, and the protein bands were visualized by immunoblotting with anti–His_6_ antibody (Thermofisher, Cat No. MA1–21315–A488).

#### Single molecule DNA curtain assays

All single molecule measurements were carried out at 30°C. Experiments were conducted with a custom–built prism–based total internal reflection fluorescence (TIRF) microscope (Nikon) equipped with 488–nm laser (Coherent Sapphire, 200 mW) and a 561–nm laser (Coherent Sapphire, 200 mW) (De Tullio et al., 2018). To prevent spectral overlap, GFP and mCherry images were collected using 500 ms alternating shuttering of the two lasers, and data was collected using 100–ms integration time with two Andor iXON X3 EMCCD cameras (De Tullio et al., 2018). Images were collected every 10 s at 50 mW laser power unless stated otherwise. All images were exported as TIFF stacks using Nikon NIS Elements software and kymographs for individual ssDNA filaments were generated and analyzed using Fiji software.

Slides were constructed by deposition of chrome barriers on quartz microscope slides via e–beam lithography and thermal evaporation as previously described (De Tullio et al., 2018; Ma et al., 2017b). Briefly, holes were drilled into a quartz microscope slide with a diamond coated drill bit. The slides were then cleaned for 20 minutes in piranha solution (3 parts H_2_SO_4_ to 1 part H_2_O_2_). Followed by washing 3x with H_2_O. The slide surfaces were then coated first with Poly–methyl methacrylate (PMMA) 25 K (Polymer Source, Cat. No. P9790–MMA), then PMMA 495K (MicroChem, Cat# M130003), and finally aquaSave (Mitsubishi Rayon Co., Ltd., Cat# aquaSAVE–53za). Patterns were written into the PMMA using e–beam lithography. Slides were developed in an isopropanol: methyl–isobutyl ketone solution (3:1) with sonication for 1 min at 0°C. Chromium was then deposited on the microscope surface using electron beam evaporation. The PMMA and excess chromium were then removed by lift–off in acetone. The quality of the chromium features was checked by light microscopy. Flow cells were assembled and dsDNA curtains were prepared as previously described (Greene et al., 2010). First, using double–sided tape a glass coverslip was attached to the quartz microscope slide. The tape was melted in an oven to create a sealed reaction chamber. Inlet and outlet nanoports were then glued over the holes that had been drilled in the microscope slide and the flow cell was completed. ssDNA was generated using a 5’ biotinylated primer annealed to M13 circular single–stranded DNA as a template, by rolling circle replication with phi29 DNA polymerase (Ma et al., 2017a). The flow cell was attached to the microfluidic system and passivated with streptavidin containing lipid bilayer as described (De Tullio et al., 2018; Ma et al., 2017b). Single–stranded DNA (ssDNA) was injected into the flow cell, attached to the lipid bilayer through biotin–streptavidin linkage, aligned at the barriers by application of a constant buffer flow, and secondary structures were disrupted by addition of 7 M urea in BSA buffer (40 mM Tris–Cl [pH 8.0], 2 mM MgCl_2_, 1 mM DTT, and 0.3 mg mL^−1^ BSA) at a flow rate of 0.8 mL min^−1^. Free ssDNA ends were anchored to the exposed pedestals through non–specific adsorption.

#### Rad51 filament assembly and kinetics

RPA filaments were assembled using BSA buffer (40 mM Tris–Cl [pH 8.0], 2 mM MgCl_2_, 1 mM DTT, and 0.3 mg mL^−1^ BSA) supplemented with 100 pM unlabeled RPA, GFP–RPA, or mCherry– RPA (as indicated) for at least 8 min at a flow rate of 0.8 mL min^−1^. Rad51 filament assembly was initiated by injecting 2 μM Rad51 (or 5 μM Rad51–K191R, as indicated) with or without Rad55– Rad57 in HR buffer (30 mM Tris–Ac [pH 7.5], 50 mM KCl, 20 mM MgCl2, 2 mM ATP (or 2 mM AMP–PNP, as indicated), 1 mM DTT, 0.2 mg mL^−1^ BSA), followed by a 20 min incubation in the absence of buffer flow. At the end of this incubation, unbound proteins were washed away by applying HR buffer at a flow rate of 0.8 mL min^−1^ for at least 1 min.

#### In vitro photobleaching controls and Hed1–binding assays

2 μM Rad51 with 60 nM GFP–Rad55–Rad57 was co–injected in HR buffer (30 mM Tris–Ac [pH 7.5], 50 mM KCl, 20 mM MgCl2, 2 mM ATP, 1 mM DTT, 0.2 mg mL^−1^ BSA), onto mCherry– RPA–ssDNA filaments. Data was collected for 15 mins under stopped flow conditions, and images acquired either every 10 s, 20 s, or 60 s. Mean normalized GFP intensity and 95% CI was plotted as a function of time for each frame rate. For experiments with Hed1, 2 μM Rad51 with 30 nM GFP–Hed1 was co–injected in HR buffer onto mCherry–RPA–ssDNA filaments. Images were acquired every 10 s during a 15 min incubation under stopped flow conditions and mean normalized intensity with 95% CI plotted.

#### Srs2 antirecombinase assays

Sample chambers containing pre–assembled Rad51–ssDNA filaments were flushed with HR buffer for several minutes to remove any unbound Rad51 prior to the Srs2 injections. 500 pM Srs2 was injected in HR buffer supplemented with 500 pM RPA and observed under stopped flow conditions. In experiments with free Rad51 and/or free Rad55–Rad57 in the buffer, the indicated amount of each protein was co–injected with Srs2 and RPA in 150 μL aliquots using a flow rate of 0.5 mL min^−1^, and then observed under stopped flow conditions.

### QUANTIFICATION AND STATISTICAL ANALYSIS

#### Single–molecule data collection and analysis

For all two–color images, we used a custom–built shuttering system to avoid signal bleed–through during image acquisition (De Tullio et al., 2018). With this system, images from the green (GFP) and the red (mCherry) channels are recorded independently, these recordings are offset by 100 ms such that when one camera records the red channel image, the green laser is shuttered off, and vice versa. This system prevents any possible signal bleed–through between the two channels. Images were captured at an acquisition rate of 1 frame per 10 s (unless indicated otherwise) with a 100– millisecond integration time, and the laser was shuttered between each acquired image to minimize photo–bleaching. Raw TIFF images were imported as image stacks into ImageJ, and kymographs were generated from the image stacks by defining a 1–pixel wide region of interest (ROI) along the long–axis of the individual dsDNA molecules. All data analysis was performed using the resulting kymographs.

#### Rad51 assembly kinetics

For calculating assembly kinetics, kymographs were generated by defining a 1–pixel–wide region of interest (ROI) along the length of individual ssDNA–filaments. Integrated signal intensity for each time point was obtained from individual kymographs and normalized using the maximum value for that kymograph. The mean and 95% CI of normalized intensities across all ssDNA molecules was plotted for each time point, and the distribution was fit to a single exponential decay curve using GraphPad Prism to obtain the rate constants.

#### Analysis of residual Rad55–Rad57

Images were acquired every 10 s with 50 mW laser power for 15 min under stopped flow conditions, after which unbound proteins were washed away by application of HR buffer. The 488 nm laser power was increased to 200 mW to visualize any residual GFP–Rad55–Rad57 remaining on Rad51–ssDNA filaments. Number of discrete GFP foci per ssDNA molecule was counted and plotted as a histogram.

#### Analysis of Srs2 trajectories

Raw TIFF files were imported as image stacks into Fiji software for analysis, and kymographs were generated by defining a 1–pixel–wide region of interest (ROI) along the length of individual ssDNA molecules. Within each kymograph, the start and end points of individual Srs2 traces were analyzed to obtain the x (time), y (position) coordinates in terms of pixel values. Apparent Srs2 translocation velocity was calculated from the number of pixels translocated per unit time = (y2 – y1)/(x2 – x1). As described previously (De Tullio et al., 2018), each pixel in the y–axis corresponds to ~725 nucleotides (nt) of the Rad51–ssDNA filament, and each pixel in the y–axis corresponds to 10s. Using these conversion factors Srs2 velocity was expressed in units of nt/s. To obtain the number of Srs2 molecules per Rad51 filament (Figure 6C,F), a single point on the Rad51–ssDNA filament was observed over a 2.5 min time window (this time window was chosen due to substantial ssDNA breakage at longer times). On each kymograph an ROI 1–pixel wide and 15– pixels long was defined parallel to the x–axis, starting from the first Srs2 trajectory. Number of Srs2 molecules per Rad51–ssDNA filament was determined by number of peaks in the GFP intensity profile across the ROI in individual kymographs (Figure S7A).

#### Statistical information

All statistical analysis was carried out using Graphpad Prism. Rates for Rad51 filament assembly were calculated by fitting the data to a single–exponential decay curve (Ma et al., 2017b). Statistical significance between groups was calculated using One–way ANOVA, with Tukey’s correction for multiple comparisons. ns = not significant, *p<0.05, **p<0.01, ***p<0.001, ****p<0.0001.

## SUPPLEMENTAL FIGURE LEGENDS

**Figure S1.**
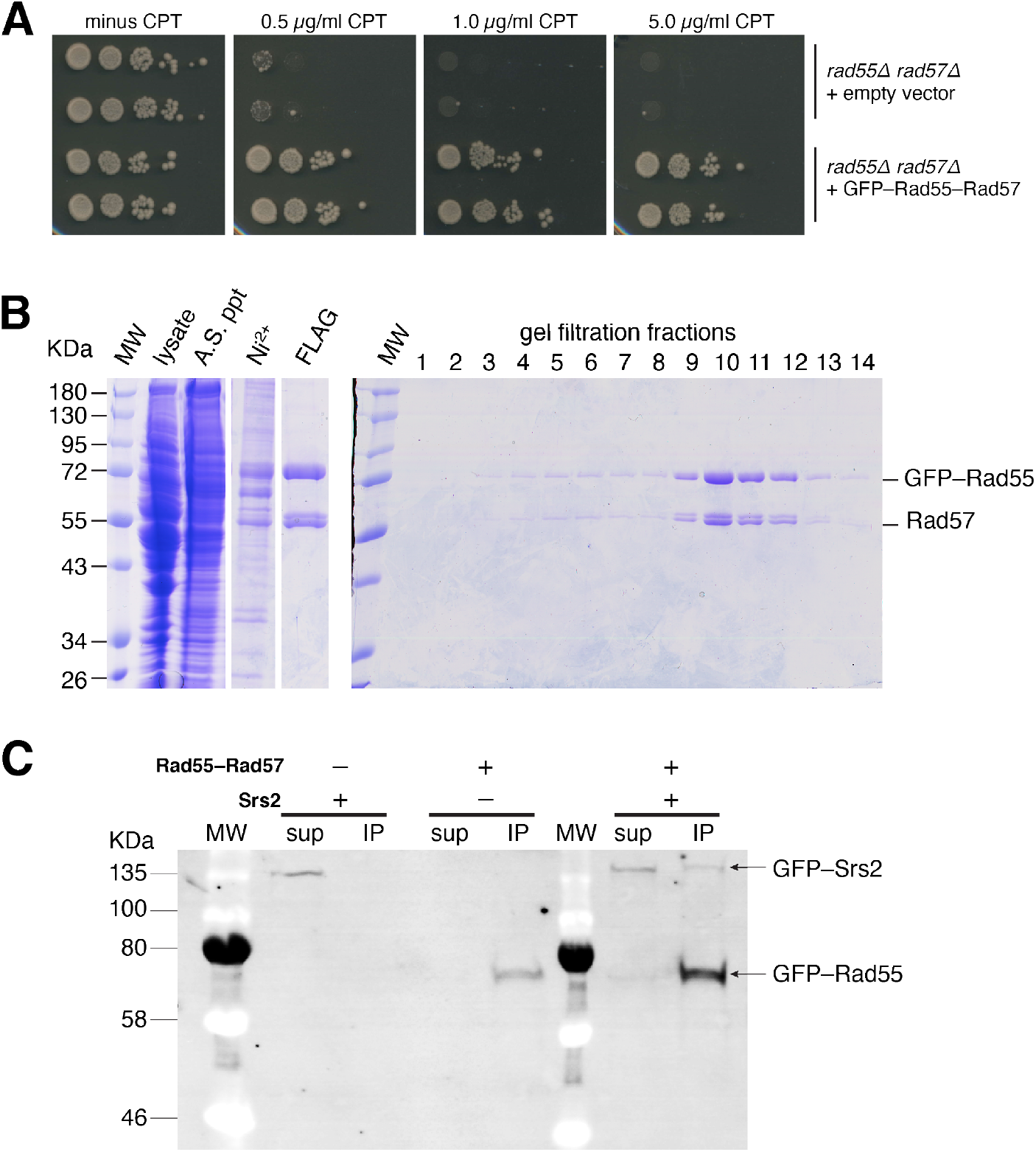
Preparation of the GFP–Rad55–Rad57 complex, related to Figure 1. (**A**) *rad55Δ. rad57Δ* cells were either complemented with empty vector, or a plasmid expressing GFP–tagged Rad55–Rad57. Colony growth of 2 separate clones was measured at the indicated concentrations of camptothecin (CPT). (**B**) Cell lysate, resuspended ammonium sulphate precipitate (A.S. ppt), elution from Ni^2+^ column, elution from FLAG column and fractions from the gel filtration column (0.5 mL fractions) were loaded on a 10% SDS PAGE gel and stained with Coomassie blue. (**C**) Pull–down with FLAG–Rad57 showing interaction of Rad55–Rad57 with GFP–Srs2. The supernatant (Sup) or bound (IP) fraction was blotted with anti–His antibody.

**Figure S2.**
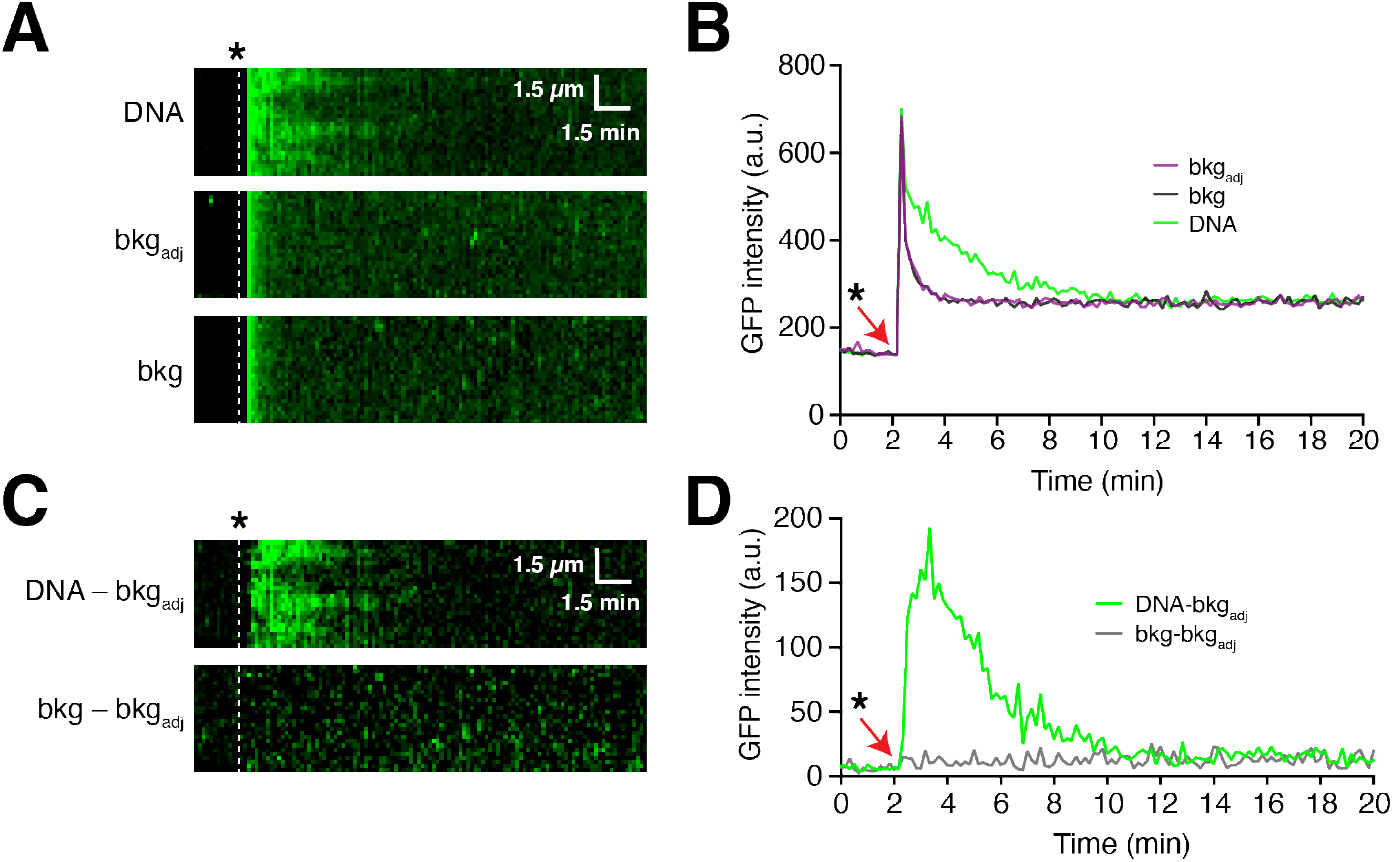
Rad55–Rad57 is depleted from mature Rad51 filaments, related to Figure 2. (**A**) Kymograph of GFP signal on a ssDNA molecule (DNA), background region adjacent to the ssDNA (bkgadj) or a distal background region away from the DNA (bkg) during co–injection of 2 μM Rad51 and 60 nM GFP–Rad55– Rad57. (**B**) Integrated GFP intensity from kymographs in (A). The time of injection of Rad51 and GFP–Rad55–Rad57 is depicted by the white dashed line or red arrow. (**C**) Resultant images after background subtraction, and (**D**) the corresponding quantification.

**Figure S3.**
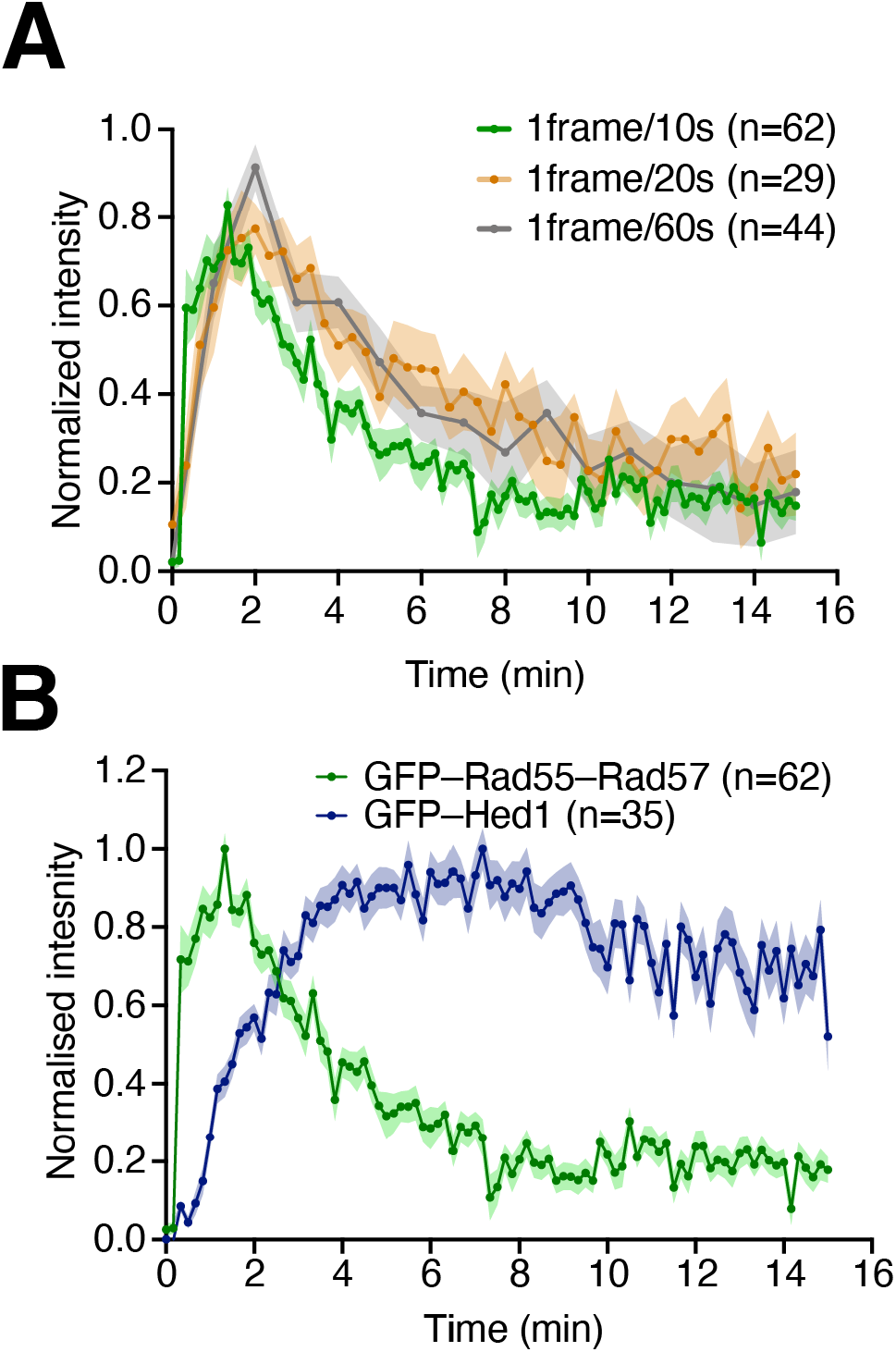
Photobleaching controls for DNA curtain assays, related to Figure 2. (**A**) Photobleaching control to confirm transient binding of Rad55–Rad57 during Rad51–ssDNA filament formation. Graph showing kinetics of Rad55–Rad57 after co–injection with Rad51 at varying laser exposure times. Normalized intensity is plotted for Rad55–Rad57 imaged at a rate of one frame per 10 s, 20 s or 60 s, as indicated; note, samples are only exposed to laser light during frame acquisition. Shaded areas represent 95% CI. (**B**) Comparison of the transient binding of GFP–Rad55–57 to the stable binding of GFP–Hed1 during Rad51–ssDNA filament formation; Hed1 is a meiosis–specific protein that binds tightly to Rad51 (Brown and Bishop, 2014; Crickard et al., 2018). Normalized intensity is plotted and the shaded area represents 95% CI.

**Figure S4.**
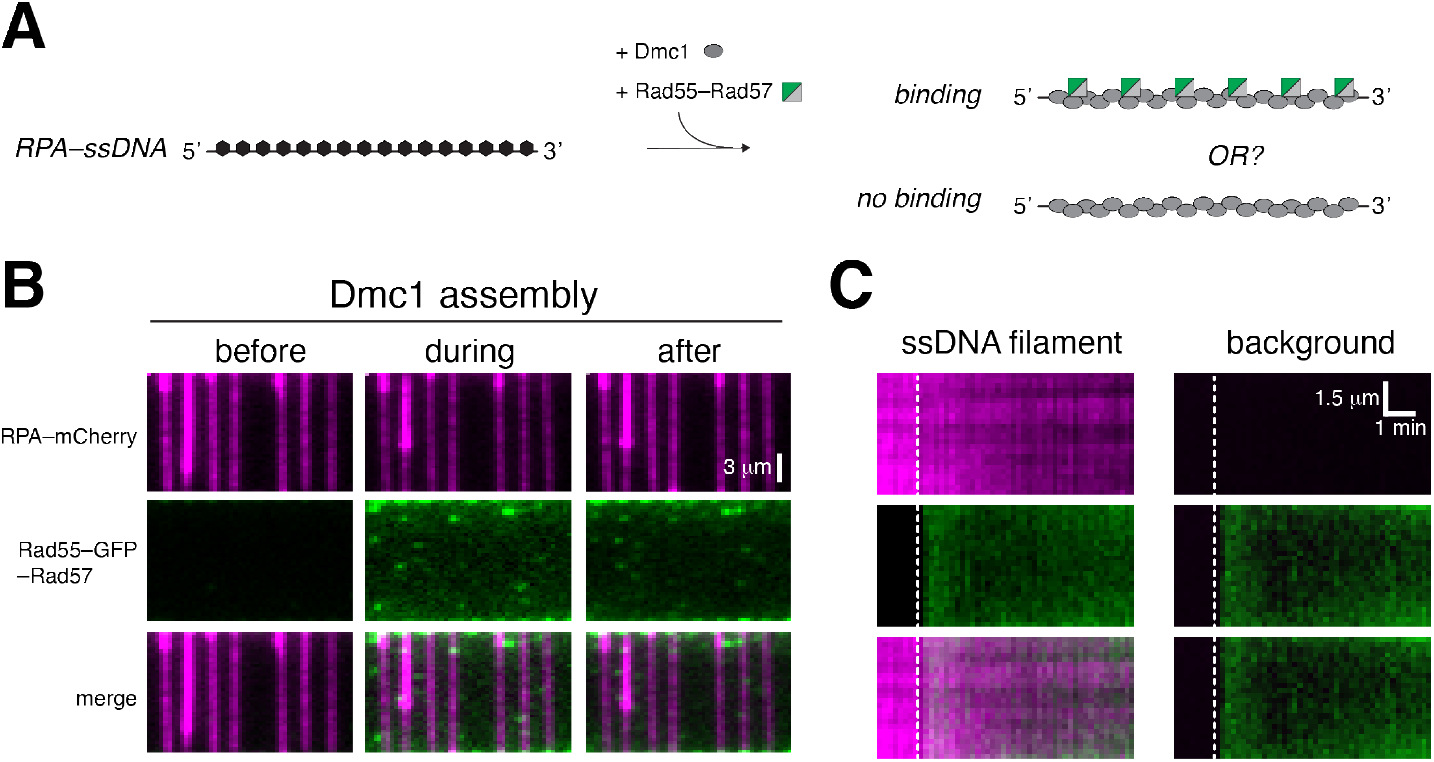
Rad55–Rad57 doesn’t bind Dmc1–ssDNA filaments, related to Figure 2. (**A**) Assay schematic to test Rad55–57 binding during Dmc1–ssDNA filament assembly. (**B**) Wide–field view of RPA–mCherry bound ssDNA (magenta) 1 min before, during, and 1 min after co–injection of 60 nM Rad55–Rad57 (green) and 2 μM Dmc1. (**C**) Representative kymographs of Dmc1–ssDNA, and background region lacking ssDNA. Injection of GFP–Rad55–Rad57 is indicated by the white dashed line.

**Figure S5.**
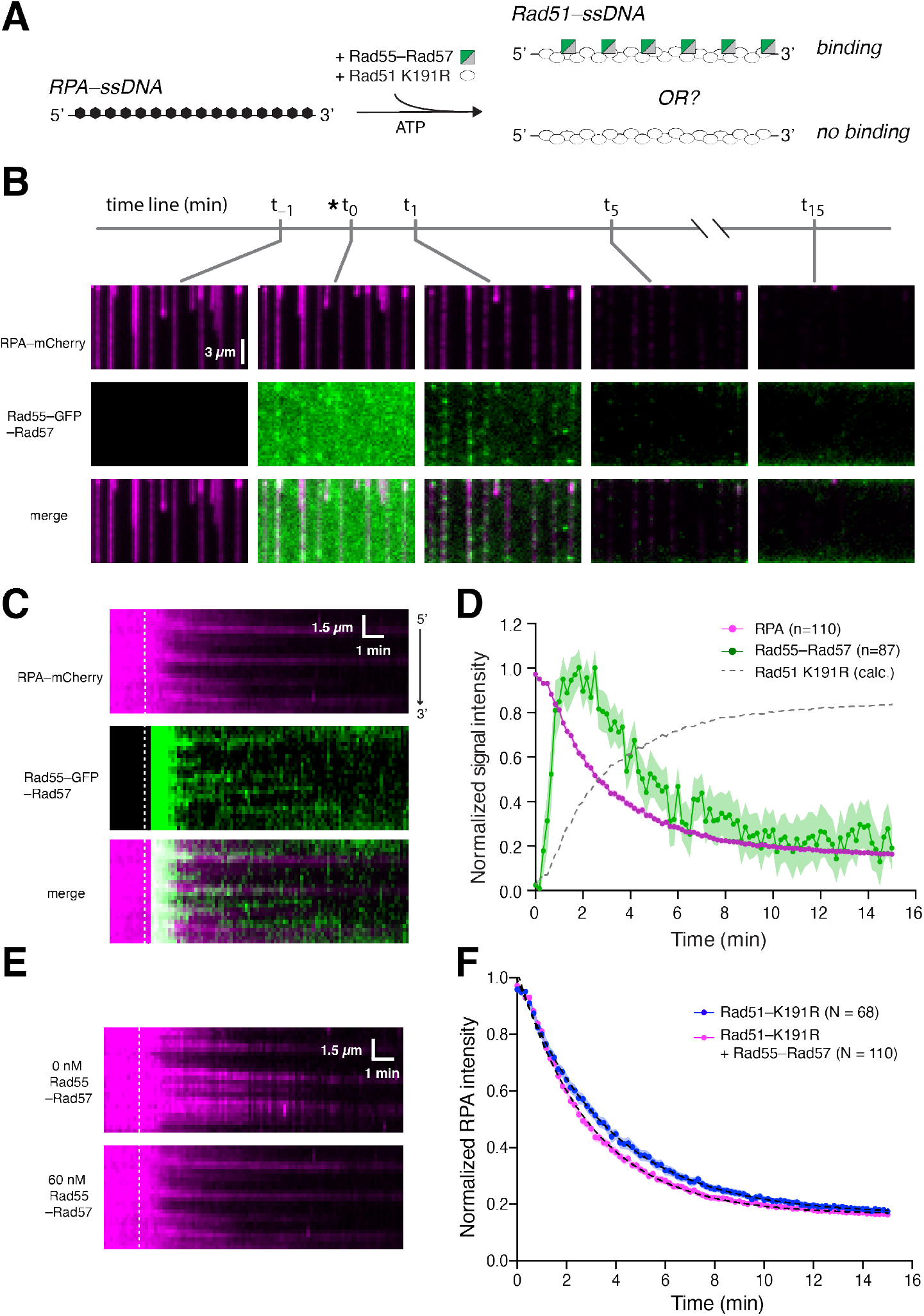
ATP hydrolysis by Rad51 is not required for binding and dissociation of Rad55– Rad57, related to Figure 4. **(A)** Experimental schematic for assays with Rad51–K191R. **(B)** Wide–field view of RPA– mCherry (magenta) bound ssDNA at the indicated time points. Time at which Rad51–K191R and Rad55–Rad57 (green) were co–injected is indicated with an asterisk (*). **(C)** Kymographs showing the behavior of mCherry–RPA and GFP–Rad55–Rad57. **(D)** Graph showing the mean normalized signal intensities for RPA–mCherry and GFP–Rad55–Rad57. The shaded area represents the 95% CI. Calculated Rad51–K191R binding is plotted as [1–RPA] signal for comparison. **(E)** Kymographs showing RPA–mCherry loss at the indicated concentrations of Rad55–Rad57 co–injected with Rad51–K191R. **(F)** Graph showing kinetics of RPA in the presence of Rad51–K191R and either, 0 nM (blue), or 60 nM Rad55–Rad57 (magenta). It should be noted that due to its reduced ssDNA affinity, we could not use lower concentrations of Rad51– K191R at which any effect of Rad55–Rad57 would be apparent. Here we use 5 μM Rad51–K191R (compared to 2 μM WT Rad51) to allow for sufficient filament assembly (Kaniecki et al., 2017).

**Figure S6.**
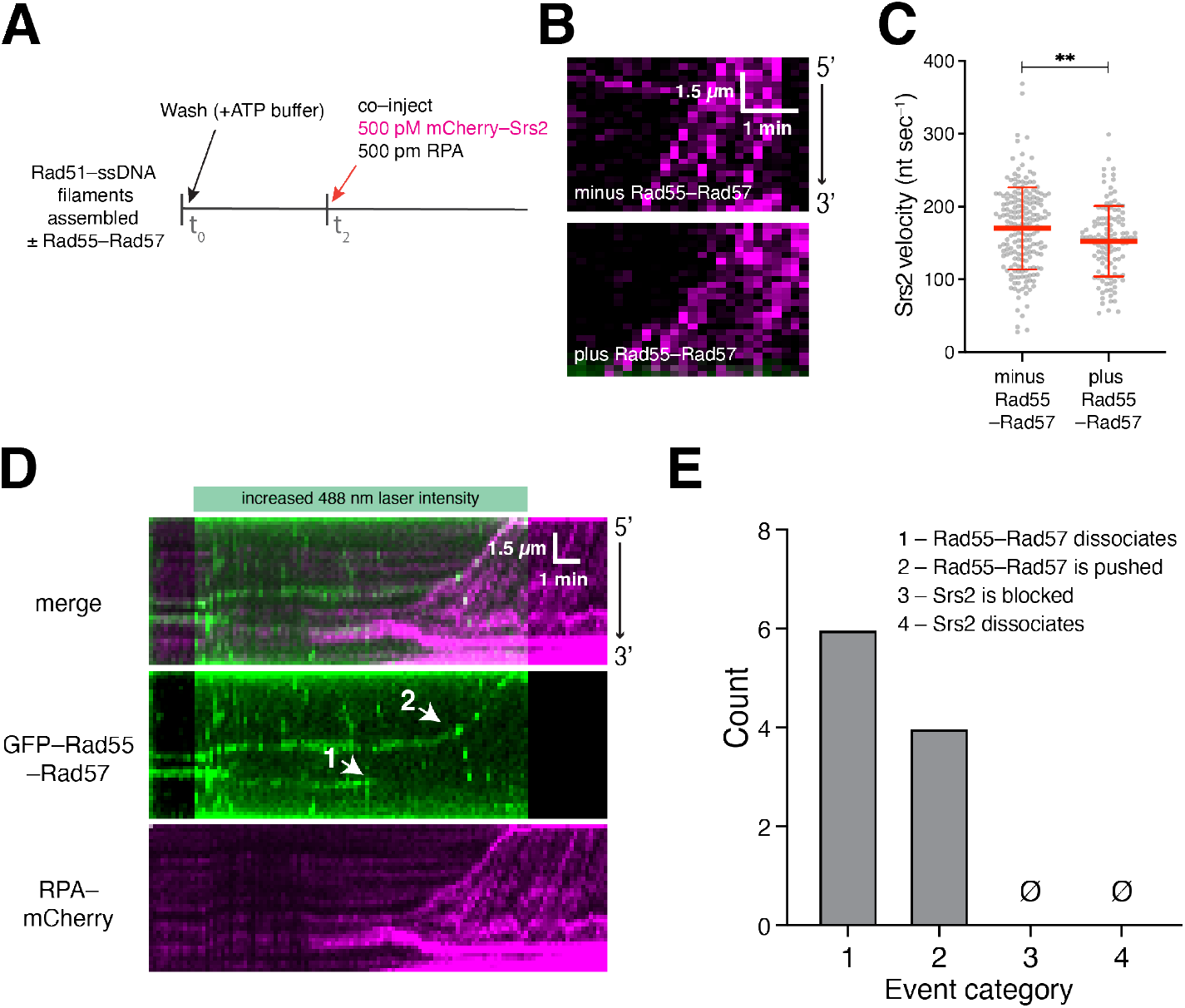
Srs2 strips residual Rad55–Rad57 from Rad51–ssDNA, related to Figure 6. **(A)** Assay schematic. (**B)** Kymographs showing mCherry–Srs2 (magenta) translocating on Rad51– ssDNA filaments formed in the absence (top) or presence (bottom) of Rad55–Rad57. **(C)** Quantification of Srs2 velocities in experiment (B). **(D)** Kymograph showing residual Rad55–57 foci (green) on Rad51–ssDNA and rebinding of RPA–mCherry (magenta) behind Srs2. Translocating Srs2 stripped off Rad55–Rad57 foci or pushed it along the ssDNA in a 3’→5’ direction (indicated by white arrows). **(E)** Quantification of observed outcomes when Srs2 encountered Rad55–Rad57 foci on DNA.

**Figure S7.**
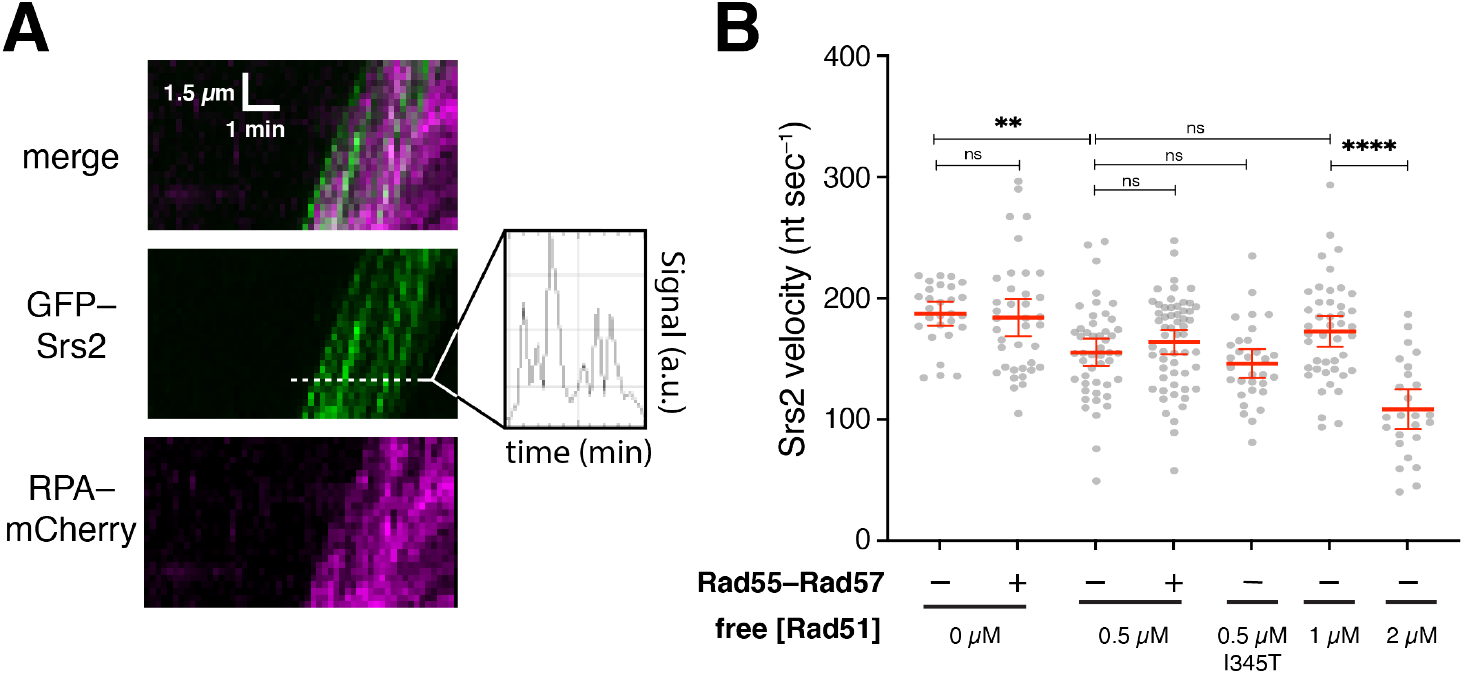
Quantification of GFP–Srs2 activity in the presence of free Rad51, RPA and Rad55–Rad57, related to Figure 6. **(A)** Quantifications were done over a 2.5 min time window (as indicated) starting from the first Srs2 molecule translocation on Rad51–ssDNA filaments. Number of Srs2 molecules per Rad51– ssDNA filament was determined by number of peaks in the GFP intensity profile from individual kymographs. **(B)** Quantification of Srs2 velocities with 0.5 nM free RPA–mCherry, ± 60 nM Rad55–Rad57 and 0–2 μM Rad51 or 0.5 μM Rad51–I345T, as indicated. The red lines indicate the means ± 95% CI for each data set.

**Table S1.** Rad51 filament assembly parameters, related to Figures 2, 3, 4 & S5.

|  | Filament assembly conditions | Fluorescence signal | Dissociation rate<br>k (min<sup>-1</sup>) | 95% CI | N | Fig. |
| --- | --- | --- | --- | --- | --- | --- |
| **1** | 2 μM Rad51 + 2 mM ATP | RPA–mCherry | 0.187 | 0.181 – 0.193 | 72 | 3C |
| **2** | 2 μM Rad51 + 5 nM Rad55–Rad57 + 2 mM ATP | RPA–mCherry | 0.355 | 0.351 – 0.359 | 90 | 3C |
| **3** | 2 μM Rad51 + 60 nM Rad55–Rad57 + 2 mM ATP | RPA–mCherry | 0.493 | 0.485 – 0.501 | 59 | 3C, 2D |
|  |  | GFP–Rad55–Rad57 | 0.455 | 0.423 – 0.488 | 62 | 2D |
| **4** | 2 μM Rad51 + 2 mM AMP–PNP | RPA–mCherry | 0.129 | 0.119 – 0.140 | 80 | 4F |
| **5** | 2 μM Rad51 + 60 nM Rad55–Rad57 + 2 mM AMP–PNP | RPA–mCherry | 0.252 | 0.239 – 0.265 | 102 | 4D, 4F |
|  |  | GFP–Rad55–Rad57 | 0.020 |  | 104 | 4D |
| **6** | 5 μM Rad51–K191R + 2 mM ATP | RPA–mCherry | 0.275 | 0.270 – 0.281 | 68 | S5F |
| **7** | 5 μM Rad51–K191R + 60 nM Rad55–Rad57 + 2 mM ATP | RPA–mCherry | 0.339 | 0.334 – 0.344 | 110 | S5D,F |
|  |  | GFP–Rad55–Rad57 | 0.348 | 0.305 – 0.393 | 87 | S5D |

**Table S2.** Srs2 translocation parameters, related to Figures 6, S6 & S7.

|  | Reaction description | Srs2 co-injected with | Srs2 velocity (nt sec<sup>-1</sup>) |  |  | No. of Srs2 molecules |  |  | Fig. |
| --- | --- | --- | --- | --- | --- | --- | --- | --- | --- |
|  |  |  | mean | s.d. | N | mean | s.d. | N |  |
| **1** | 500 pM GFP–Srs2 on pre-assembled Rad51 filaments | 500 pM RPA–mCherry | 175.3 | ±33.2 | 55 | 6.6 | ±1.4 | 47 | 6B,C, S7B |
| **2** | 500 pM GFP–Srs2 on pre-assembled Rad51 filaments | 500 pM RPA–mCherry | 163.6 | ±48.8 | 65 | 6.5 | ±1.2 | 64 | 6E,F, S7B |
|  |  | 60 nM GFP–Rad55–Rad57 |  |  |  |  |  |  |  |
| **3** | 500 pM GFP–Srs2 on pre-assembled Rad51 filaments | 500 pM RPA–mCherry | 156.2 | ±36.8 | 59 | 6.4 | ±1.4 | 61 | 6B–F, S7B |
|  |  | 0.5 µM Rad51 |  |  |  |  |  |  |  |
| **4** | 500 pM GFP–Srs2 on pre-assembled Rad51 filaments | 500 pM RPA–mCherry | 169.2 | ±41.6 | 44 | 3.9 | ±1.7 | 34 | 6B,C, S7B |
|  |  | 1 µM Rad51 |  |  |  |  |  |  |  |
| **5** | 500 pM GFP–Srs2 on pre-assembled Rad51 filaments | 500 pM RPA–mCherry | 105.1 | ±39.8 | 25 | 2.5 | ±1.5 | 26 | 6B,C, S7B |
|  |  | 2 µM Rad51 |  |  |  |  |  |  |  |
| **6** | 500 pM GFP–Srs2 on pre-assembled Rad51 filaments | 500 pM RPA–mCherry | 153.9 | ±35.61 | 63 | 4.5 | ±1.8 | 68 | 6E,F, S7B |
|  |  | 0.5 µM Rad51 |  |  |  |  |  |  |  |
|  |  | 60 nM Rad55–Rad57 |  |  |  |  |  |  |  |
| **7** | 500 pM GFP–Srs2 on pre-assembled Rad51 filaments | 500 pM RPA–mCherry | 142.7 | ±32.61 | 31 | 4.9 | ±1.7 | 36 | 6E,F, S7B |
|  |  | 0.5 µM Rad51–I345T |  |  |  |  |  |  |  |
| **8** | 500 pM mCherry–Srs2 on pre-assembled Rad51 filaments | 500 pM unlabeled RPA | 170 | ±56.5 | 194 | n.a. |  |  | S6B,C |
| **9** | 500 pM mCherry–Srs2 on Rad51 filaments pre-assembled with 60 nM Rad55–Rad57 | 500 pM unlabeled RPA | 152 | ±48.5 | 119 | n.a. |  |  | S6B,C |

